# Data Processing Choices Can Affect Findings in Differential Methylation Analyses: An Investigation Using Data from the LIMIT RCT

**DOI:** 10.1101/2022.06.14.496049

**Authors:** Jennie Louise, Andrea R Deussen, Jodie M Dodd

**Affiliations:** Discipline of Obstetrics & Gynaecology and The Robinson Research Institute, The University of Adelaide, Adelaide, South Australia, AUSTRALIA; Adelaide Health Technology Asseessment, The University of Adelaide, Adelaide, South Australia, AUSTRALIA; Department of Perinatal Medicine, Women’s and Babies Division, The Women’s and Children’s Hospital, Adelaide, South Australia, AUSTRALIA

## Abstract

**Objective:** A wide array of methods exist for processing and analysing DNA methylation data. We aimed to perform a systematic comparison of the behaviour of these methods, using cord blood DNAm from the LIMIT RCT, in relation to detecting hypothesised effects of interest (intervention and pre-pregnancy maternal BMI) as well as effects known to be spurious, and known to be present.

**Methods:** DNAm data, from 645 cord blood samples analysed using Illumina 450K BeadChip arrays, were normalised using three different methods (with probe filtering undertaken pre- or post-normalisation). Batch effects were handled with a supervised algorithm, an unsupervised algorithm, or adjustment in the analysis model. Analysis was undertaken with and without adjustment for estimated cell type proportions. The effects estimated included intervention and BMI (effects of interest in the original study), infant sex and randomly assigned groups. Data processing and analysis methods were compared in relation to number and identity of differentially methylated probes, rankings of probes by p value and log-fold-change, and distributions of p values and log-fold-change estimates.

**Results:** There were differences corresponding to each of the processing and analysis choices. Importantly, some combinations of data processing choices resulted in a substantial number of spurious ‘significant’ findings. We recommend greater emphasis on replication and greater use of sensitivity analyses.

**Clinical Trials Registration:** ACTRN12607000161426

## Introduction and Background

With the advent of high-throughput assays, epigenome-wide DNA methylation studies have become more popular, and researchers are now investigating the effects on DNA methylation (DNAm) of a wide range of environmental exposures and physiological conditions, with particular interest in the contribution of epigenetic mechanisms such as DNAm to the early life origins of health and disease. The ability to perform EWAS is particularly useful in relation to conditions where associated differences in DNAm are likely to be fairly modest (1). However, DNAm data – as with high-dimensional ‘omics’ data generally – requires substantial pre-processing prior to analysis, including probe and sample filtering, normalisation to remove variation due to technological factors, and correction for other factors which may confound effects of interest, such as batch effects or differences in cell type proportions between samples. Numerous methods exist to perform these processing steps, but there is no clear consensus on the best processing or analysis approach (2,3).

We recently investigated the effect of an antenatal diet and lifestyle intervention, and of maternal early pregnancy BMI, on neonatal cord blood DNA methylation in infants of mothers who were overweight or obese in early pregnancy. The findings are reported in a companion paper. In brief, we did not find evidence of differential methylation in relation to either the intervention or early pregnancy maternal BMI, and were unable to replicate findings from previous studies which reported a range of loci to be significantly differentially methylated in relation to these factors. Moreover, in conducting sensitivity analyses involving use of different normalisation methods and methods for handling batch effects, we observed a number of differences in results, both in relation to the most highly ranked probes and in the ‘detection’ of effects which we believed to be spurious. We therefore set out to investigate the impact of different data-processing choices in a more systematic way, looking at the effect of these choices on detection of differentially methylated probes (DMPs), on overall distribution of p values and log-fold-change (logFC) estimates, and on rankings of probes (by logFC and by p value).

In this paper we report the findings from a set of analyses conducted on data processed and handled according to a combination of different prespecified choices. The particular factors we investigated related to (a) filtering of probes before versus after normalisation; (b) method used for normalisation; (c) method used to handle batch effects; and (d) adjustment vs non-adjustment for estimated cell type proportions. We compared results from analyses estimating effects of the antenatal intervention and maternal early pregnancy BMI, but also were interested in observing the behaviour of different data-processing choices in relation to effects that were both known to be present, and known to be absent. We therefore performed further analyses in which effects of infant sex, and of randomly assigned (fake) groups were estimated.

## Data and Methods

### The LIMIT Randomised Controlled Trial

The LIMIT study was a randomised, controlled trial of an antenatal diet and lifestyle intervention for women with early pregnancy BMI ≥25.0 kg/m^2^. The study, and its primary and main secondary outcomes, have been extensively reported elsewhere (4). Women were eligible if they had early pregnancy BMI ≥25.0 kg/m^2^, a singleton pregnancy between 10^+0^ and 20^+0^ weeks’ gestation, and no previously existing diabetes. A total of 2212 women were randomised to receive either Lifestyle Advice (n=1108), a comprehensive diet and lifestyle intervention, or Standard Care (n=1104), in which antenatal care was delivered according to local guidelines (and did not include information on diet or physical activity). The primary outcome was birth of a Large for Gestational Age (LGA) infant. There were no significant differences observed between the groups in relation to this outcome; however, a significantly lower incidence of birthweight >4kg was observed in the Lifestyle Advice group, with a Relative Risk of 0.82 (95% CI: 0.68, 0.99, p=0.04). Additionally, measures of diet quality and physical activity were improved in women in the Lifestyle Advice group compared to those in the Standard Care group (5).

Cord Blood DNA for a range of secondary studies was collected at the time of birth from consenting participants, and was frozen as whole blood preserved with EDTA. Funding was available to perform DNA methylation analysis for a total of 649 samples, which were randomly selected from the total number of available samples, balanced between the Lifestyle Advice and Standard Care groups. After DNA extraction, genome-wide DNA methylation was performed using the Illumina Infinium HumanMethylation 450K Bead-Chip array. Results were supplied as raw probe intensities (*idats* files).

For the additional analyses investigating known spurious effects, artificial (fake) groups were created by assigning samples based on random draws from binomial distributions. The first grouping (‘Tortoiseshell’ vs’Tabby’) was generated using a binomial distribution with 50% probability of assignment to each group. The second grouping (‘Long’-vs ‘Short-Haired’) was created to mimic stratified randomisation as well as unequal proportions in each group: within each level of the first fake group, samples were assigned to Long-Haired with 40% probability and Short-Haired with 60% probability.

All data processing and analyses were undertaken using R version 4.0 (6).

### Probe and Sample Filtering

The *minfi* package (7) was used to read in the raw *idats* files (without normalisation), and to calculate both probe-wise and sample-wise ‘detection p values’. Samples were identified as ‘faulty’ if they had a detection p-value ≥0.05; 13 such samples were excluded; however these were due to a known chip failure, and had subsequently been rerun. A further four samples were excluded because the correct corresponding study identifier could not be ascertained, leaving 645 samples for analysis.

Probes were filtered using multiple criteria. Firstly, probes were excluded if they had a detection p-value ≥0.001 in more than 25% of the 645 samples, indicating that their signal could not be accurately detected for a large proportion of samples (8,9). Secondly, probes were excluded if they were on a list of those previously identified as cross-reactive (10); i.e. there was a high probability they may hybridize to locations on the genome different to those for which the probe was designed (8,11). Thirdly, probes with an identified SNP within 3 nucleotides of the CpG site and minor allele frequency >1%, and probes on the X and Y chromosomes were excluded. This was done in order to avoid spurious methylation ‘differences’ due either to SNPs within the CpG targets, or due to X and Y chromosomes (8,11). Filtering of cross-reactive probes, probes with a nearby SNP, and probes on the X and Y chromosomes, was performed using the *DMRCate* package (12). This left 426,572 probes available for analysis.

Probe filtering was performed either after normalisation (post-filtered) or prior to normalisation (pre-filtered). The one exception for pre-filtering was when normalising using the BMIQ method, where probes on the X and Y chromosome were retained as this was required in order for the function to run.

### Normalisation

Normalisation involves making changes to the raw data in order to remove artifactual variation. In the case of Illumina 450K BeadChip arrays, this requires correcting for the presence of two different probe types. Infinium I probes use the same colour signal for methylated and unmethylated CpG and are often used for regions of high CpG density, while Infinium II probes use different colours to differentiate between methylated and unmethylated states (13,14). In general, the distribution of *β* values 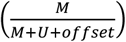 will be bimodal, with peaks corresponding to methylated and unmethylated states), but the distribution of Infinium II probes differs from that of Infinium I, being more compressed towards 0.5 (13) and hence having a smaller ‘dynamic range’ (8,15).

Numerous different methods exist for normalising Illumina BeadChip array data, but there is little consensus or guidance on which should be employed in a given context. The main advice is that ‘between-array’ methods, which normalise across samples, are preferable when global differences between samples are expected, while ‘within-array’ methods, which normalise probes within each sample, are better suited to effects in which the majority of genes will not be differentially expressed. (9) The latter is the context in which many EWAS studies, including the present one, are conducted; as noted above, only modest differences in a small proportion of genes are expected for most early-life exposures. The methods chosen for the present investigation have all been used in the context of studies such as this: Categorical-Subset Quantile Normalisation (SQN) (14,16), Beta-Mixture Quantile Normalisation (BMIQ) (15), and Subset-Quantile Within-Array Normalisation (SWAN) (18). While numerous other methods exist, a comparison of all available normalisation methods was beyond the scope of this paper. Further details on the methods are given in the Supplementary Information.

Both Subset Quantile Normalisation and Subset-Within-Array-Normalisation were performed using functions in the *minfi* package (*preprocessQuantile* and *preprocessSWAN* respectively), on raw intensity data. Beta-Mixture Quantile normalisation was performed using the *champ*.*norm* function in the *ChAMP* package after converting intensities to *β* values.

### Batch Effects

Batch effects arise when samples are processed in separate groups, creating unwanted variation due, for example, to different reagents, different plates or different scanner settings. (3,19,20) There are 12 Illumina 450K arrays (samples) per chip (this is reduced to 8 arrays per chip for the more recent 850K array); thus most studies involving large numbers of samples must be run on multiple chips. This introduces extra variability to the data, and may also confound the actual effects of interest, if samples from different groups are not evenly distributed between the batches. These effects must be accounted for in order to obtain valid estimates of the effects of interest.

Unlike probe filtering and normalisation, batch effects can be handled at the analysis stage, by adjusting for batch in the analysis model. However, it is also common to address batch effects at the data-processing stage, using a batch-correction algorithm, with the resulting data considered to be free of batch effects (19). The ComBat algorithm has been widely used and considered the most effective method (2) of removing batch effects in DNAm data; it has been incorporated into various analysis pipelines. Until recently, ComBat could be implemented only as a supervised method, in which the biological factors of interest had to be specified along with the batch variable (3) (21); it can now also be implemented as an unsupervised method, in which only the batch variable is specified.

For each of the normalised datasets (i.e. SQN, BMIQ and SWAN normalised datasets, each with probes filtered either before or after normalisation), we handled batch effects in three ways: firstly, by adjusting for batch in the analysis model (BatchAdjust); secondly, implementing the supervised ComBat algorithm (sCB); and thirdly, implementing the unsupervised ComBat algorithm (uCB). For the supervised ComBat algorithm, it was necessary to run the process twice: once with the effects of interest specified as maternal early pregnancy BMI, antenatal intervention group, and their interaction; and again with the effects of interest specified as Fake Group 1, Fake Group 2, their interaction, and infant sex.

### Cell Type Proportions

Cord blood, like whole blood, contains a mixture of different cell types, which have different DNA methylation profiles.(22,23) If samples differ in the proportion of these different cell types, this may confound effects of interest, either hiding true differences in DNAm, or giving rise to spurious differences. Most studies of the effect of BMI, lifestyle interventions, or similar factors on cord blood DNA methylation have not attempted (or have not documented an attempt) to correct for potential differences in cell type composition, perhaps because reference profiles for cord blood were not available until more recently (24), and the mix of cell types and DNAm profiles may differ in cord blood compared to whole blood, making it inappropriate to apply reference profiles from whole blood to cord blood data.(25)

We estimated the proportion of B cells, CD4+T, CD8+T, granulocytes, monocytes, natural killer, and nucleated RBCs in the raw data using the *estimateCellCounts()* function in the *minfi* package, with the Cord Blood reference panel. The estimated proportions were then added to the metadata for use as adjustment variables in the analyses. We then undertook analyses either adjusted or not adjusted for estimated cell type proportion.

Figure 1 depicts the combinations of data-processing and analysis choices that were undertaken. In brief, there were six normalised datasets (three different normalisation methods, with probe filtering performed before normalisation or after normalisation). These datasets were either used immediately for analysis, or processed using the ComBat algorithm (in both supervised and unsupervised form) prior to analysis. Non-ComBat-processed data were analysed with three different models: an unadjusted model (containing only the effects of interest), a model adjusted for batch, and a model adjusted for batch and estimated cell type proportions. ComBat-processed data were analysed with two different models: one containing no other adjustment variables (but assumed to be ‘pre-adjusted’ for batch), and one adjusted for cell type proportion.

**Figure 1.**
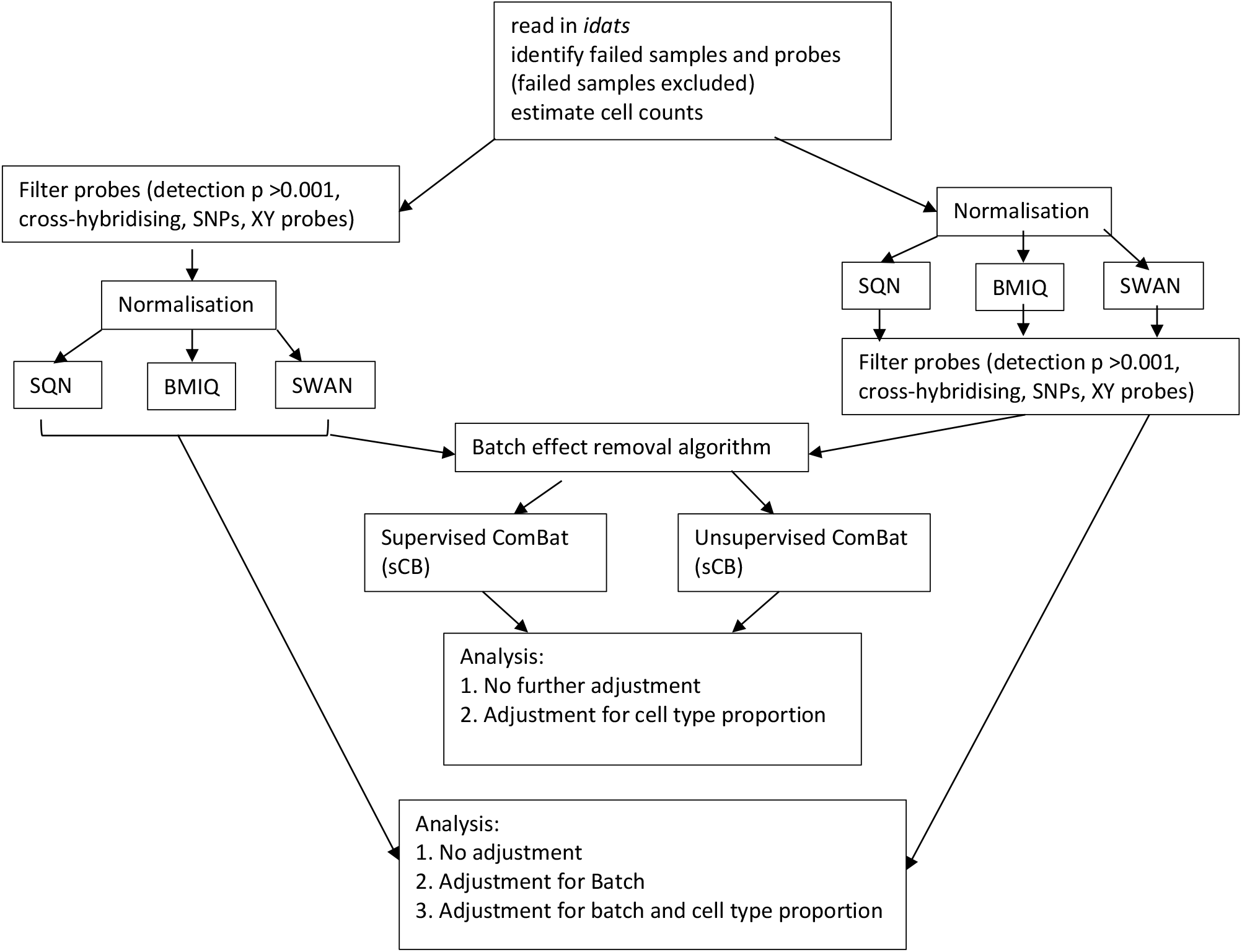
Flowchart of Data Processing and Analysis

### Statistical Analysis

Differential methylation was investigated probe-wise using linear models with empirical Bayes variance correction as implemented in the *limma* package (26,27). For effects of BMI and intervention, models specified BMI (as continuous and mean-centred), intervention (Lifestyle Advice vs Standard Care), and their interaction. Contrasts were specified for the effect of the intervention at the mean BMI of the cohort, and at 5 kg/m^2^ above the mean; and for the effect of an increase of 5 kg/m^2^ in BMI in each of the intervention groups (Standard Care, Lifestyle Advice). For effects of fake groups and infant sex, the models specified sex (Female vs Male), Fake Group 1 (Tortoiseshell vs Tabby), Fake Group 2 (Long-Haired vs Short-Haired), and their interaction. Contrasts were specified for infant sex, and for the effect of each fake group separately within levels of the other fake group (i.e. effect of Tortoiseshell in Long-Haired and in Short-Haired; and effect of Short-Haired in Tortoiseshell and Tabby).

For each contrast in each model, the number and identity (where applicable) of any differentially methylated probes (DMPs) were obtained. For detection of DMPs, *limma*’s default method of multiple-comparisons correction (Benjamini-Hochberg) was used; this method controls the False Discovery Rate, or the proportion of statistically significant results not corresponding to true effects. Where DMPs were obtained, a comparison was made using the Holm method, which controls the Family-Wise-Error Rate (the probability that at least one statistically significant result does not correspond to a true effect). The full set of p values and estimated log-fold-changes (for all 426572 probes) corresponding to each contrast were also obtained, in order to compare probe rankings and overall distributions. To make the comparison more tractable, probe rankings were investigated using only those probes ranked in the top 10 (i.e. the probes with the smallest p value, or largest estimated logFC, in a given model).

The findings from different data-processing choices were then compared along five dimensions:

1. Number and identity of differentially methylated probes (DMPs); for infant sex, the direction of differential methylation (‘down’, corresponding to negative t-statistics or lower methylation in females, versus ‘up’, corresponding to higher methylation in females) was also examined.
2. Consistency of rankings by p value for ‘top 10’ probes;
3. Consistency of rankings by logFC for ‘top 10’ probes, as well as the consistency of the logFC estimates;
4. Overall distribution of p values;
5. Overall distribution of logFC estimates.

## Results

All data processing choices had an impact on downstream analysis results, in terms of the number (and identity) of differentially methylated probes, rankings of probes (by p value and logFC), estimates of logFC, and overall distribution of p values and logFC, corresponding to both real and spurious effects of interest. In some cases a consistent impact of a particular choice was observed, while in others there was no consistent pattern, or this pattern varied according to the other choices with which it was combined.

Tables 1-3 give information about differentially methylated probes in each of the models fitted for the combinations of filtering, normalisation, batch correction and cell adjustment approaches. Table 1 lists the number of significant negative (‘down’) probes and significant positive (‘up’) probes for infant sex. Table 2 gives the number of differentially methylated probes for any of the BMI and intervention effects, while Table 3 gives the number of differentially methylated probes for any of the fake group effects (the specific effects and probes, as well as their rankings in other models, are listed in Supplementary Tables 3-8).

**Table 1.** Number of DMPs for Infant Sex (Female), by Probe Filtering Method, Batch Correction Method, Normalisation Method and Cell Type Method.

| Model |  | SQN |  | BMIQ |  | SWAN |  |
| --- | --- | --- | --- | --- | --- | --- | --- |
|  |  | Post Filtered | Pre Filtered | Post Filtered | Pre Filtered | Post Filtered | Pre Filtered |
| No ComBat |  |  |  |  |  |  |  |
| - Unadjusted | Down | 15088 | 14878 | 7935 | 7554 | 7581 | 6890 |
|  | Up | 20587 | 20441 | 28004 | 30741 | 25784 | 25962 |
| - Adjusted for Batch | Down | 13132 | 13215 | 6602 | 6112 | 7100 | 6140 |
|  | Up | 20225 | 19956 | 39482 | 34408 | 30362 | 29780 |
| - Adjusted for Batch + Cell | Down | 28406 | 28633 | 15855 | 14900 | 14239 | 10408 |
|  | Up | 31973 | 32204 | 39719 | 45255 | 43155 | 40709 |
| Supervised ComBat |  |  |  |  |  |  |  |
| - Unadjusted* | Down | 20967 | 21230 | 11235 | 10180 | 10772 | 9252 |
|  | Up | 25690 | 25320 | 41518 | 48191 | 45193 | 43237 |
| - Adjusted for Cell | Down | 35559 | 36022 | 18036 | 16512 | 16423 | 12198 |
|  | Up | 37068 | 36972 | 56521 | 64851 | 68769 | 65109 |
| UnSupervised ComBat |  |  |  |  |  |  |  |
| - Unadjusted* | Down | 14603 | 14763 | 7634 | 6836 | 7344 | 6377 |
|  | Up | 21336 | 21041 | 30961 | 34882 | 31892 | 31170 |
| - Adjusted for Cell | Down | 28012 | 28286 | 14060 | 12916 | 13030 | 9560 |
|  | Up | 32478 | 32447 | 43370 | 49520 | 52037 | 49102 |

\* No adjustment beyond the correction for batch as implemented in the ComBat algorithm

**Table 2.** DMPs for Intervention and BMI Effects.

| Model | SQN |  | BMIQ |  | SWAN |  |
| --- | --- | --- | --- | --- | --- | --- |
|  | Post Filtered | Pre Filtered | Post Filtered | Pre Filtered | Post Filtered | Pre Filtered |
| No ComBat |  |  |  |  |  |  |
| - Unadjusted | 0 | 0 | 7 | 11 | 0 | 0 |
| - Adjusted for Batch | 0 | 0 | 8 | 6 | 0 | 0 |
| - Adjusted for Batch + Cell | 0 | 0 | 6 | 8 | 0 | 0 |
| Supervised ComBat |  |  |  |  |  |  |
| - Unadjusted | 1 | 1 | 0 | 99 | 0 | 0 |
| - Adjusted for Cell | 3 | 3 | 9 | 208 | 6 | 6 |

| UnSupervised ComBat |  |  |  |  |  |  |
| --- | --- | --- | --- | --- | --- | --- |
| - Unadjusted | 0 | 0 | 0 | 0 | 0 | 0 |
| - Adjusted for Cell | 0 | 0 | 0 | 0 | 0 | 0 |

**Table 3.** DMPs for Fake Groups.

|  | SQN |  | BMIQ |  | SWAN |  |
| --- | --- | --- | --- | --- | --- | --- |
| Model | Post Filtered | Pre Filtered | Post Filtered | Pre Filtered | Post Filtered | Pre Filtered |
| No ComBat |  |  |  |  |  |  |
| - Unadjusted | 0 | 0 | 0 | 0 | 0 | 0 |
| - Adjusted for Batch | 2180 | 2574 | 0 | 0 | 0 | 0 |
| - Adjusted for Batch + Cell | 0 | 0 | 0 | 0 | 0 | 0 |
| Supervised ComBat |  |  |  |  |  |  |
| - Unadjusted | 6768 | 7007 | 3 | 6 | 8 | 8 |
| - Adjusted for Cell | 124 | 134 | 1 | 2 | 0 | 0 |
| UnSupervised ComBat |  |  |  |  |  |  |
| - Unadjusted | 0 | 0 | 0 | 0 | 0 | 0 |
| - Adjusted for Cell | 0 | 0 | 0 | 0 | 0 | 0 |

Differences in ranking of probes by p value and log-Fold-Change are shown in Figures 2 and 3, respectively, for a selected set of effects (infant sex; the effect of maternal BMI in the Standard Care group, and the effect of ‘short-haired’ in ‘Tabby’). Similarly, the overall distribution of p values and of log-Fold-Change estimates for this same set of selected effects is shown in Figures 4 and 5.

**Figure 2.**
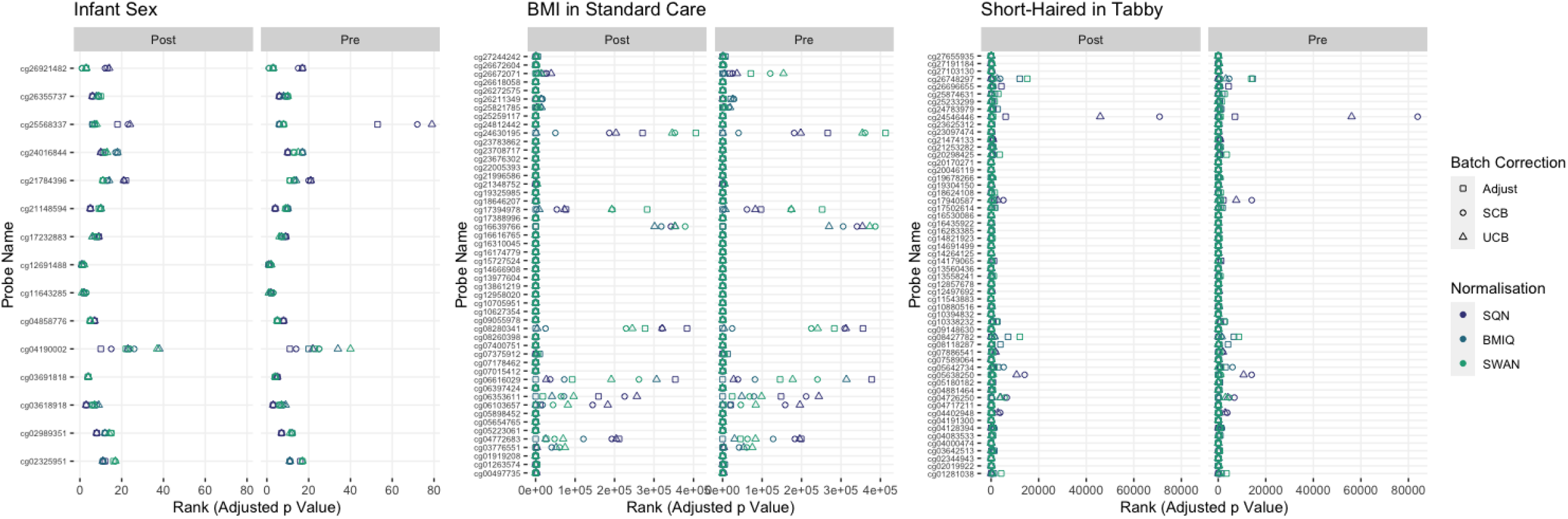
Probes Ranked in Top 10 by P value in Batch+Cell Adjusted Model, for (a) Infant Sex, (b) BMI in Standard Care, (c) Short-Haired in Tabby. For each probe, the rank is given by pre- vs post-filtering, normalisation method, and batch-handling method. The model is one adjusting for batch (either explicitly in the model or via batch-correction algorithm) and cell type proportion.

**Figure 3.**
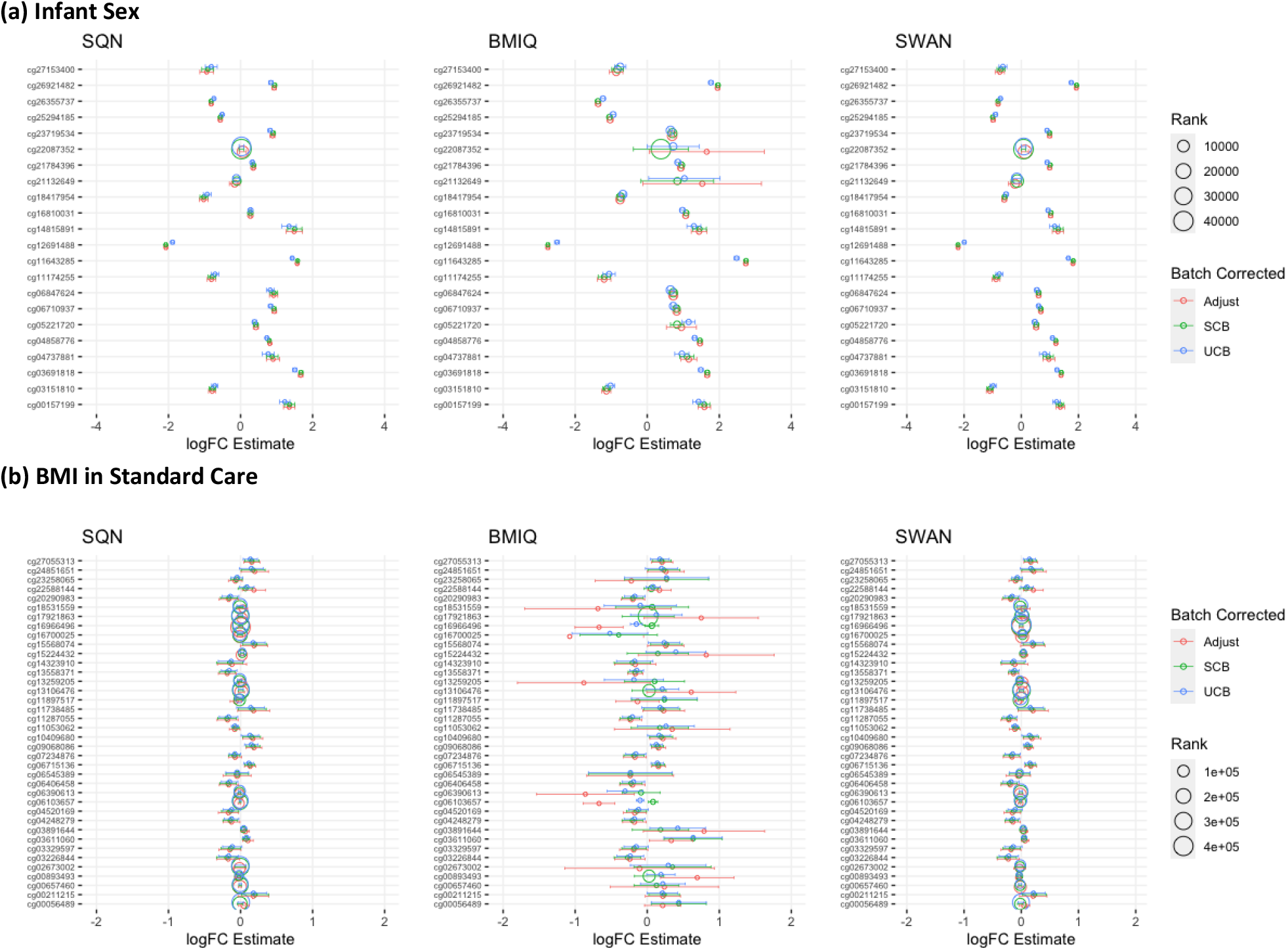
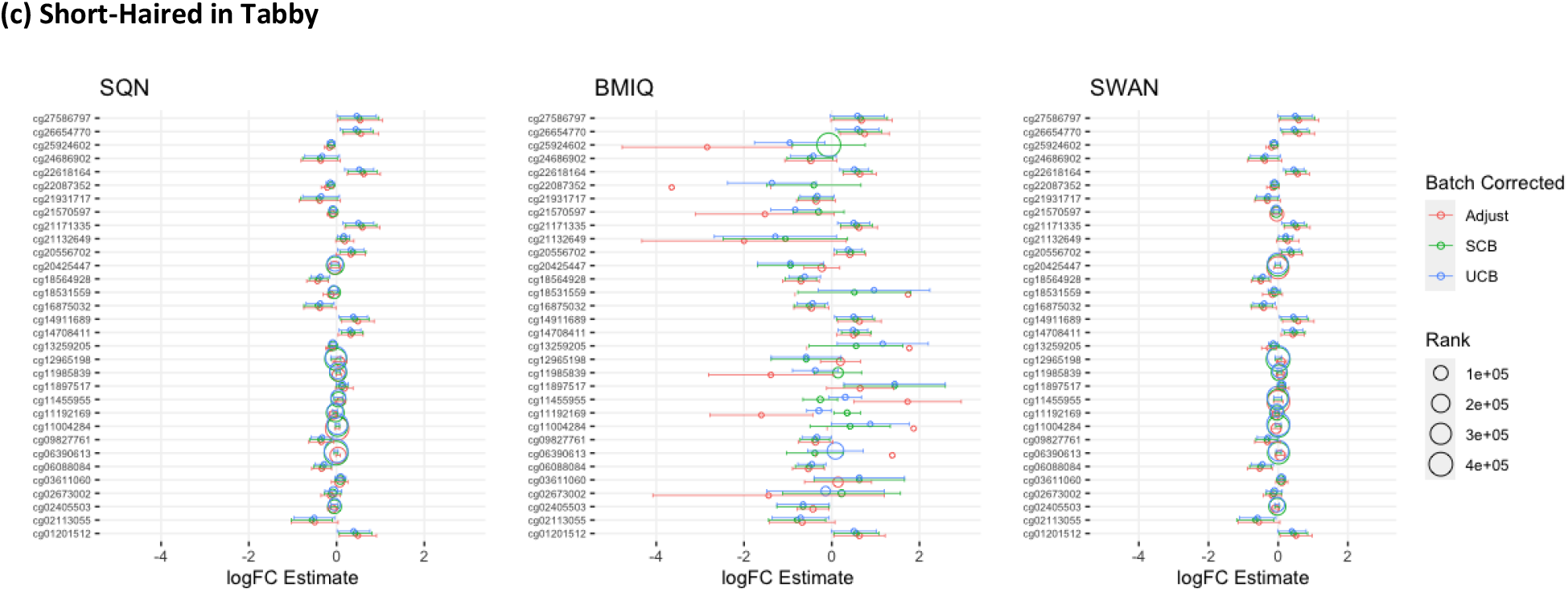
Probes Ranked in Top 10 in any Batch and Cell-Adjusted Model by Log Fold Change, for (a) Infant Sex, (b) BMI in Standard Care, (c) Short-Haired in Tabby. Only models from data with probe filtered post-normalisation are included, to simplify results presentation. The model is one adjusting for batch (either explicitly in the model or via batch-correction algorithm) and cell type proportion. The graphs give the estimated log-Fold-Change (circles) and 95% confidence interval by normalisation method and batch-correction method, with rank also indicated by the size of the circles.

**Figure 4.**
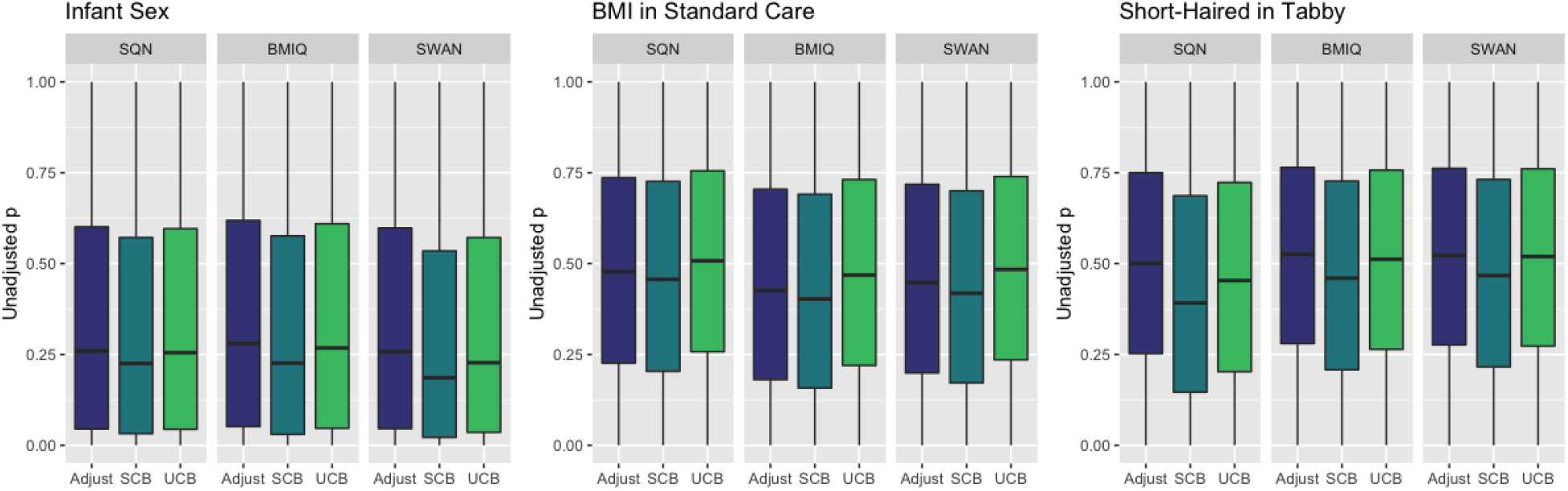
Distribution of Unadjusted P Values by Normalisation and Batch Correction Method, for Batch and Cell Adjusted Models. Only models from data where probe filtering was performed post-normalisation are included. The model is one adjusting for batch (either explicitly in the model or via batch-correction algorithm) and cell type proportion.

**Figure 5.**
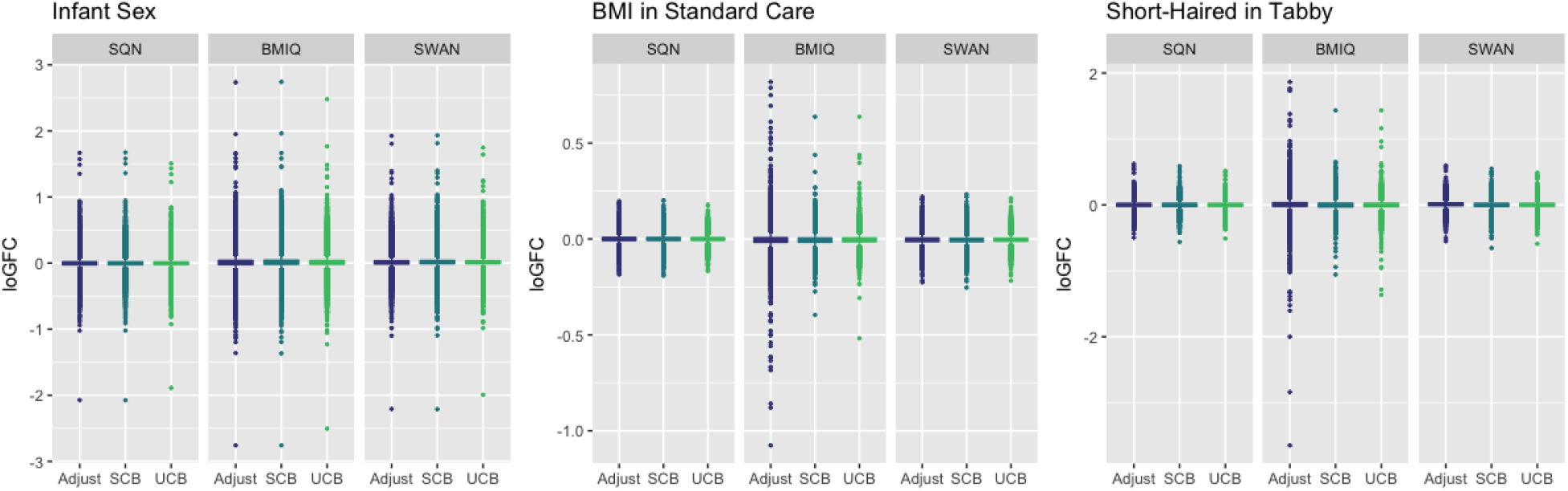
Distribution of Log-Fold-Change Estimates by Normalisation and Batch Correction Method, for Batch and Cell Adjusted Models. Only models from data where probe filtering was performed post-normalisation are included. The model is one adjusting for batch (either explicitly in the model or via batch-correction algorithm) and cell type proportion.

### Effect of probe filtering pre-normalisation vs post-normalisation

Filtering probes prior to normalisation, compared to filtering after normalisation, led to modest differences in number of DMPs, rankings of probes by logFC and p value, and overall distributions of p values and logFC estimates. Filtering pre-normalisation produced different numbers of DMPs for infant sex, but the nature of the effect differed by normalisation method: in SWAN data there was a consistent pattern of fewer significant probes both negative and positive, while in BMIQ data there were fewer negative but more positive probes, and in SQN data there were more negative but fewer positive probes. In relation to effects of BMI, intervention, and fake groups, differences were harder to discern due to the lack of any DMPs for many models; however, when DMPs were present for an effect, there was a tendency for there to be a greater number of them in the pre-filtered data.

Probe rankings, by logFC and p value, tended to be relatively consistent between pre-filtered and post-filtered data, with some cases of larger discrepancies in rankings for individual probes. The discrepancies were more common, and larger, for fake group, BMI and intervention effects than for infant sex. Similarly, there were no dramatic differences in distributions of p values or logFC estimates for infant sex; there were differences in distribution between pre- and post-filtered data for fake group, intervention and BMI effects, but there was no consistent pattern to these differences.

### Effect of Normalisation Method

Normalisation method had a substantial influence on number and identity of DMPs, rankings of probes and p values, and distributions of p values and logFC estimates. For infant sex, SQN data consistently had the highest number of significant negative probes and the lowest number of significant positive probes, while SWAN data always had the lowest number of significant positive probes. For BMI and intervention effects, only BMIQ data produced DMPs where no ComBat processing was used; in data processed using supervised ComBat, all three normalisation methods resulted in some DMPs, but the number and identity of these probes differed. In fake group data, SQN data produced a large number of significant probes in non-ComBat-processed and supervised-ComBat data, while BMIQ and SWAN data produced a small number of probes in supervised-ComBat data only; again, the number and identity of the probes differed between the normalisation methods.

There was a fair degree of consistency in rankings of probes by p value for infant sex, but some large discrepancies in rankings for BMI, intervention, and fake group effects. The rankings were less consistent for highest-ranked probes by logFC, with some quite large differences in both rankings and effect estimates (including different directions of effect) for infant sex, BMI, intervention and fake groups. BMIQ estimates tended to be more extreme (further from 0) than the other two methods.

Distributions of p values and logFC estimates also differed between normalisation methods. For p valuesthe differences were not consistent across models and effects, but for logFC there was a clear difference between BMIQ and the other two methods, with the range of estimates in BMIQ data being much more widely dispersed; SQN and SWAN data had more similar distributions, but SQN was moderately narrower than SWAN across all effects and models.

### Effect of Batch Correction Method

There were clear differences in all dimensions between batch correction methods. For all effects (infant sex, BMI, intervention and fake groups), supervised ComBat processing produced a larger number of DMPs compared to either unsupervised ComBat processing or adjustment for batch in the analysis model. The difference between unsupervised ComBat and batch-adjustment was less consistent for infant sex effects, but for BMI, intervention and fake group effects, there were no DMPs in unsupervised ComBat models, whereas there were a few for batch-adjusted models.

Rankings of top probes by p value were relatively consistent between batch-adjustment methods for infant sex, but there were some large discrepancies particularly for BMI and intervention effects, and especially in BMIQ data. The same phenomenon was observed for logFC rankings, which also showed a tendency for logFC estimates in unsupervised-ComBat data to be smaller in absolute magnitude (closer to 0).

The distribution of p values showed clear and consistent differences between batch-correction methods, with the distribution in supervised ComBat data shifted down substantially relative to both unsupervised ComBat and batch-adjusted models, for all effects. For logFC estimates, supervised ComBat and batch-adjusted data were generally fairly similar, but unsupervised ComBat data generally resulted in a narrower range.

### Effect of Adjustment for Estimated Cell Type Proportion

Adjustment for cell type proportion produced different results, but the impact differed depending on the effect. Adjustment for batch resulted in a substantially larger number of DMPs (both negative and positive) for infant sex, but reduced the number of DMPs for fake groups (for models where there were DMPs for fake groups effects). For BMI and intervention, the effect of cell type adjustment was mostly but not entirely to produce more DMPs.

The effect of cell type adjustment on top probe rankings was fairly modest, although some quite large discrepancies were observed for p value rankings, logFC rankings, and logFC estimates. The effect on distribution of p values depended on the effect: for infant sex, adjustment for cell type proportion consistently (for all normalisation and batch-correction methods) shifted the distribution downwards, whereas the differences were less consistent and smaller in BMI, intervention and fake group effects. There were no large or consistent differences in distribution of logFC estimates between cell-type-adjusted and non-adjusted models.

## Discussion

Different choices in probe filtering, normalisation, batch handling, and adjustment for cell types resulted in different findings regarding the presence and identity of differentially methylated probes, rankings of probes by p value and log-fold-change, and different overall distributions of p values and log-fold-change estimates. Some differences were relatively modest, while others were more substantial. The effect of some choices appeared to be consistent: in particular, supervised ComBat processing resulted in more differentially methylated probes and shifted the distribution of p values downward, while unsupervised ComBat processing resulted in a narrower range of log-fold-change estimates which were generally closer to 0 (a null effect) than other methods. Data normalised using BMIQ tended to produce a more widely-dispersed distribution of log-fold-change estimates and also appeared to respond in a more volatile manner to other data processing choices. By contrast, the effect of filtering probes prior to normalisation, and of adjustment for estimated cell type proportion, had different impacts depending on other data processing choices, and the effects being studied. Of interest, the number of differentially methylated probes for infant sex was increased, and the p value distribution shifted downward, when models were adjusted for estimated cell type proportion, suggesting that the removal of noise due to cell type differences allowed more precise estimation of these effects. However, the effect in relation to fake group effects was equivocal.

The results of these analyses are consistent with other investigations which have been undertaken into different data-processing and analysis choices. In particular, the potential for ‘false positives’ to result from supervised batch-correction methods specifying effects of interest has been previously identified by a number of authors.(2,3,19). The finding that the distribution of p values in the supervised ComBat algorithm tends to shift the p value distribution downward is consistent with the finding of Nygaard et al that, in contexts where the effects of interest are not evenly spread between batches, the distribution of F-statistics will be biased upwards (19). While implementation as an unsupervised method may be preferable, our findings suggest that this may create a different problem, with the estimates of log-fold-change corresponding to effects of interest biased towards zero.

In terms of different normalisation approaches, Wu et al (17) compared a variety of normalisation approaches, including GenomeStudio, SWAN, BMIQ, and a ‘complete pipeline’ incorporating SQN, when investigating the association between smoking and cord blood methylation. They found that with more stringent Type I error control, and for the “most confident” results, the different normalisation methods gave similar values; more differences arose with laxer Type I error control. When using a split-data method to validate findings, many “significant” differences at the CpG level in the development data did not validate in the testing data. They noted a tendency for more statistically significant differences to arise in SQN data, which they hypothesise may be due to reduced overall variance. In our investigation, the main context in which SQN data produced a large number of spurious differentially methylated probes was when supervised ComBat, or adjustment for batch in the model, was used. This suggests that batch adjustment is particularly ill-advised in the context of SQN normalisation; since SQN involves between-array as well as within-array normalisation, additional adjustment for batch may be over-correcting.

Our findings do not suggest that there is one particular combination of methods which can be guaranteed to ‘work’ in all contexts, although there are some recommendations which can be made. Echoing Nygaard et al, we suggest that adjustment for batch in the model, rather than batch-correction algorithms, be used. Secondly, as many other authors have noted, researchers working with DNAm data should better understand the methods built into standard pipelines (2,3), and should better document the specific data-processing methods used (2,19). The use of a more stringent method of Type I error control may also help to reduce the number of spurious findings: the use of FDR correction methods such as Benjamini-Hochberg, while very common (9), may not be sufficient to deal with higher rates of spurious results (19). In our data, the use of the Holm method (which controls the Family-Wise Error Rate) reduced, but did not eliminate, spurious findings associated with fake group effects. Investigation of DNA regions, rather than probe-wise analysis, may also help to differentiate true methylation differences from spurious ones (14).

It is also important, in our view, to pay more attention to the context in which a particular epigenome-wide analysis is performed. For example, a less stringent method of Type I error control may often be chosen because the study is exploratory (hypothesis-generating) rather than confirmatory, and it is considered more important not to miss potential findings than to rule out spurious ones. In this case, the results from such studies should be interpreted accordingly: as suggestive findings which cannot be confidently accepted until they are validated in new data. The validation of existing findings should be treated as a high priority in epigenetics research (3).The degree of confidence that can be placed in any new discoveries could be enhanced by performing sensitivity analyses – re-performing analyses using different normalisation methods, batch correction methods, or models - which we believe should become standard in this area.

## Notes

### Competing Interest Statement

The authors have declared no competing interest.

## References

1. Marabita F, Almgren M, Lindholm ME, Ruhrmann S, Fagerström-Billai F, Jagodic M, et al. An evaluation of analysis pipelines for DNA methylation profiling using the Illumina HumanMethylation450 BeadChip platform. Epigenetics. 2013 Mar 1;8(3):333–46.

2. Zindler T, Frieling H, Neyazi A, Bleich S, Friedel E. Simulating ComBat: how batch correction can lead to the systematic introduction of false positive results in DNA methylation microarray studies. BMC Bioinformatics. 2020 Jun 30;21(1):271.

3. Price EM, Robinson WP. Adjusting for Batch Effects in DNA Methylation Microarray Data, a Lesson Learned. Front Genet [Internet]. 2018 Mar 16 [cited 2020 Sep 14];9. Available from: https://www.ncbi.nlm.nih.gov/pmc/articles/PMC5864890/

4. Dodd JM, Turnbull D, McPhee AJ, Deussen AR, Grivell RM, Yelland LN, et al. Antenatal lifestyle advice for women who are overweight or obese: LIMIT randomised trial. BMJ. 2014 Feb 10;348(feb 10 3):g1285–g1285.

5. Dodd JM, Cramp C, Sui Z, Yelland LN, Deussen AR, Grivell RM, et al. The effects of antenatal dietary and lifestyle advice for women who are overweight or obese on maternal diet and physical activity: the LIMIT randomised trial. BMC Med. 2014 Oct 13;12(1):161.

6. R Core Team. R: A Language and Environment for Statistical Computing [Internet]. Vienna, Austria: R Foundation for Statistical Computing; 2018 [cited 2020 Sep 2]. Available from: https://www.r-project.org/

7. Aryee MJ, Jaffe AE, Corrada-Bravo H, Ladd-Acosta C, Feinberg AP, Hansen KD, et al. Minfi: a flexible and comprehensive Bioconductor package for the analysis of Infinium DNA methylation microarrays. Bioinformatics. 2014 May 15;30(10):1363–9.

8. Dedeurwaerder S, Defrance M, Bizet M, Calonne E, Bontempi G, Fuks F. A comprehensive overview of Infinium HumanMethylation450 data processing. Brief Bioinform. 2014;15(6):929–41.

9. Maksimovic J, Phipson B, Oshlack A. A cross-package Bioconductor workflow for analysing methylation array data. F1000Research. 2017 Apr 5;5:1281.

10. Chen Y, Lemire M, Choufani S, Butcher DT, Grafodatskaya D, Zanke BW, et al. Discovery of cross-reactive probes and polymorphic CpGs in the Illumina Infinium HumanMethylation450 microarray. Epigenetics. 2013 Feb;8(2):203–9.

11. Naeem H, Wong NC, Chatterton Z, Hong MKH, Pedersen JS, Corcoran NM, et al. Reducing the risk of false discovery enabling identification of biologically significant genome-wide methylation status using the HumanMethylation450 array. BMC Genomics. 2014;15(1):51.

12. Peters TJ, Buckley MJ, Statham AL, Pidsley R, Samaras K, V Lord R, et al. De novo identification of differentially methylated regions in the human genome. Epigenetics Chromatin. 2015 Jan 27;8(1):6.

13. Pidsley R, CC YW, Volta M, Lunnon K, Mill J, Schalkwyk LC. A data-driven approach to preprocessing Illumina 450K methylation array data. BMC Genomics. 2013;14:293–293.

14. Wang T, Guan W, Lin J, Boutaoui N, Canino G, Luo J, et al. A systematic study of normalization methods for Infinium 450K methylation data using whole-genome bisulfite sequencing data. Epigenetics. 2015;10(7):662–9.

15. Teschendorff AE, Marabita F, Lechner M, Bartlett T, Tegner J, Gomez-Cabrero D, et al. A beta-mixture quantile normalization method for correcting probe design bias in Illumina Infinium 450 k DNA methylation data. Bioinformatics. 2013 Jan 15;29(2):189–96.

16. Wu MC, Joubert BR, Kuan P, Håberg SE, Nystad W, Peddada SD, et al. A systematic assessment of normalization approaches for the Infinium 450K methylation platform. Epigenetics. 2014 Feb 1;9(2):318–29.

17. Hicks SC, Irizarry RA. quantro: a data-driven approach to guide the choice of an appropriate normalization method. Genome Biol. 2015 Dec;16(1):117.

18. Maksimovic J, Gordon L, Oshlack A. SWAN: Subset-quantile Within Array Normalization for Illumina Infinium HumanMethylation450 BeadChips. Genome Biol. 2012 Jun 15;13(6):R44.

19. Nygaard V, Rødland EA, Hovig E. Methods that remove batch effects while retaining group differences may lead to exaggerated confidence in downstream analyses. Biostatistics. 2016 Jan 1;17(1):29–39.

20. Morris TJ, Beck S. Analysis pipelines and packages for Infinium HumanMethylation450 BeadChip (450k) data. Methods. 2015;72(C):3–8.

21. Fortin J-P, Triche T, Hansen K. Preprocessing, normalization and integration of the Illumina HumanMethylationEPIC array. 2016 Jul 23 [cited 2018 Jun 21]; Available from: http://biorxiv.org/lookup/doi/10.1101/065490

22. Jaffe AE, Irizarry RA. Accounting for cellular heterogeneity is critical in epigenome-wide association studies. Genome Biol. 2014;15(2):R31–R31.

23. Teschendorff AE, Zheng SC. Cell-type deconvolution in epigenome-wide association studies: a review and recommendations. Epigenomics. 2017 May;9(5):757–68.

24. Bakulski KM, Feinberg JI, Andrews SV, Yang J, Brown S, L McKenney S, et al. DNA methylation of cord blood cell types: Applications for mixed cell birth studies. Epigenetics. 2016 May 3;11(5):354–62.

25. Cardenas A, Allard C, Doyon M, Houseman EA, Bakulski KM, Perron P, et al. Validation of a DNA methylation reference panel for the estimation of nucleated cells types in cord blood. Epigenetics. 2016 Nov;11(11):773–9.

26. Ritchie ME, Phipson B, Wu D, Hu Y, Law CW, Shi W, et al. limma powers differential expression analyses for RNA-sequencing and microarray studies. Nucleic Acids Res. 2015;43(7):e47–e47.

27. Smyth GK. limma: Linear Models for Microarray Data. Bioinforma Comput Biol Solut Using R Bioconductor. (2005):397–420.

