## Supplementary Information for "Data Processing Choices Can Affect Findings in Differential Methylation Analyses: An Investigation Using Data from the LIMIT RCT"

### Details of Normalisation Methods Used

Most widely used normalisation methods for Illumina 450K BeadChip data all involve some form of Quantile Normalisation, in which probes within some group or category are ranked by intensity, and their values then replaced by the mean intensity of the probes with the same rank within the group . (16) (13,14). The differences between the methods are in how the groups, within which quantile normalisation is performed, are defined. , and our aim was not to exhaustively catalogue the differences between the methods, but rather to investigate whether making different choices between methods might result in different findings.

Categorical-Subset Quantile Normalisation (SQN) assumes that probes belonging to the same CpG category (e.g. CpG islands or shores) should have similar distributions (14). Probes are separated into two groups based on their relationship to CpG islands, and the quantile normalisation method is applied separately to these two groups. This is therefore a between-array method, which normalises distributions across samples (16).

SWAN, like SQN, is based on an assumption that probes (regardless of type) which have similar characteristics should have similar distributions. In this case, the relevant characteristic is the number of CpG sites on the probe, rather than relationship to CpG islands. Another difference from SQN is that SWAN is a within-array normalisation method. It forms subsets, based on the number of CpG sites in probes, and then uses quantile normalisation to ensure that the distributions within these subsets are similar (18).

BMIQ is a model-based method which does not make assumptions about the distribution of probes sharing characteristics (e.g. relationship to CpG islands). Instead, it modifies the distribution of Infinium II probes to make it more consistent with that of Infinium I probes (15). A mixture model is used to assign probes to one of three methylation states (methylated, unmethylated, partially methylated); probabilities from this model are then transformed into quantiles based on the distribution of Type I probes.

**Table S1: Baseline Characteristics of LIMIT participants with Cord Blood DNAm Data**

| Characteristic | Lifestyle Advice | Standard Care | Overall |
|---|---|---|---|
| Overall Numbers | n=325 | n=320 | n=645 |
| BMI (kg/m2): Median (IQR) | 31.40 (28.10, 36.20) | 31.45 (27.98, 36.90) | 31.40 (28.00, 36.50) |
| BMI Category: (N%) | | | |
| - 25.0-29.9 | 129 (39.69) | 130 (40.62) | 259 (40.16) |
| - 30.0-34.9 | 99 (30.46) | 86 (26.88) | 185 (28.68) |
| - 35.0-39.9 | 58 (17.85) | 55 (17.19) | 113 (17.52) |
| - ≥40.0 | 39 (12.00) | 49 (15.31) | 88 (13.64) |
| Height(cm): Mean (SD) | 165.29 (6.66) | 164.73 (6.48) | 165.01 (6.57) |
| Weight(kg): Mean (SD) | 89.81 (17.48) | 89.75 (18.65) | 89.78 (18.06) |
| Parity: N(%) | | | |
| - 0 | 141 (43.38) | 128 (40.00) | 269 (41.71) |
| - 1+ | 184 (56.62) | 192 (60.00) | 376 (58.29) |
| Age at TE: Mean (SD) | 29.28 (5.56) | 29.63 (5.24) | 29.45 (5.41) |
| Smoking: N(%) | | | |
| - Yes | 274 (84.31) | 274 (85.62) | 548 (84.96) |
| - No | 47 (14.46) | 37 (11.56) | 84 (13.02) |
| - Missing | 4 (1.23) | 9 (2.81) | 13 (2.02) |
| Ethnicity: N(%) | | | |
| - Non-Caucasian | 29 (8.92) | 29 (9.06) | 58 (8.99) |
| - Caucasian | 294 (90.46) | 291 (90.94) | 585 (90.70) |
| - Missing | 2 (0.62) | 0 (0.00) | 2 (0.31) |
| SEIFA IRSD Quintile: N(%) | | | |
| - Q1 | 107 (32.92) | 87 (27.19) | 194 (30.08) |
| - Q2 | 61 (18.77) | 83 (25.94) | 144 (22.33) |
| - Q3 | 59 (18.15) | 52 (16.25) | 111 (17.21) |
| - Q4 | 46 (14.15) | 52 (16.25) | 98 (15.19) |
| - Q5 | 52 (16.00) | 46 (14.37) | 98 (15.19) |
| Infant Sex: N(%) | | | |
| - Male | 164 (50.46) | 163 (50.94) | 327 (50.70) |
| - Female | 161 (49.54) | 157 (49.06) | 318 (49.30) |
| Study Site: N(%) | | | |
| - WCH | 135 (41.54) | 136 (42.50) | 271 (42.02) |
| - FMC | 98 (30.15) | 103 (32.19) | 201 (31.16) |
| - LMH | 92 (28.31) | 81 (25.31) | 173 (26.82) |

**Table S2: Numbers assigned to fake groups**

| | Long | Short | Total |
|---|---|---|---|
| **Tabby** | 134 | 192 | 326 (50.5) |
| **Tortoiseshell** | 126 | 193 | 319 (49.5) |
| **Total** | 260 (40.3) | 385 (59.7) | 645 |

The tables below give information about probes which were significantly differentially methylated for BMI and Intervention effects, or for fake group effects, in any of the models considered. Where an effect had a large number of differentially methylated probes, the 3 probes with the smallest p value were selected for reporting. The table gives the estimated log-Fold Change, adjusted p value (using Benjamini-Hochberg method) and rank (by p value) of the probe in each model. The model(s) in which the probe was statistically significant are indicated in bold.

#### Tables S3-S5: BMI and Intervention Effects

Note that there is no table for Intervention at +5 kg/m$^2$ BMI as there were no DMPs in any model associated with this effect.

#### Table S3: Intervention at Mean BMI

| Model | Norm. | cg20260570 (chr13 C13orf34; C13orf37) | | | cg03057840 (chr19 unnamed) | | |
|---|---|---|---|---|---|---|---|
| | | logFC (95% CI)) | adj.P.Val. | rank. | logFC (95% CI)) | adj.P.Val | rank |
| Post NoCB Unadj | SQN | -0.07 (-0.11, -0.04) | 1.00 | 4 | -0.08 (-0.12, -0.04) | 1.00 | 7 |
| Post NoCB Batch | SQN | -0.08 (-0.11, -0.04) | 1.00 | 5 | -0.08 (-0.12, -0.04) | 1.00 | 8 |
| Post NoCB Batch+Cell | SQN | -0.08 (-0.12, -0.05) | 1.00 | 2 | -0.10 (-0.13, -0.06) | 0.21 | 1 |
| Post sCB Unadj | SQN | -0.08 (-0.11, -0.04) | 0.32 | 4 | -0.08 (-0.12, -0.04) | 0.45 | 11 |
| Post sCB Cell | SQN | -0.08 (-0.11, -0.05) | 0.05 | 2 | **-0.09 (-0.13, -0.06)** | **0.04** | **1** |
| Post uCB Unadj | SQN | -0.07 (-0.10, -0.04) | 1.00 | 4 | -0.08 (-0.11, -0.04) | 1.00 | 6 |
| Post uCB Cell | SQN | -0.08 (-0.11, -0.05) | 0.33 | 2 | -0.09 (-0.12, -0.05) | 0.13 | 1 |
| Pre NoCB Unadj | SQN | -0.07 (-0.11, -0.04) | 1.00 | 4 | -0.08 (-0.12, -0.04) | 1.00 | 5 |
| Pre NoCB Batch | SQN | -0.08 (-0.11, -0.04) | 1.00 | 5 | -0.08 (-0.12, -0.04) | 1.00 | 7 |
| Pre NoCB Batch+Cell | SQN | -0.08 (-0.12, -0.05) | 1.00 | 2 | -0.10 (-0.13, -0.06) | 0.20 | 1 |
| Pre sCB Unadj | SQN | -0.08 (-0.11, -0.04) | 0.33 | 4 | -0.08 (-0.12, -0.05) | 0.46 | 9 |
| Pre sCB Cell | SQN | -0.08 (-0.11, -0.05) | 0.05 | 2 | **-0.09 (-0.13, -0.06)** | **0.04** | **1** |
| Pre uCB Unadj | SQN | -0.07 (-0.10, -0.04) | 1.00 | 4 | -0.08 (-0.11, -0.04) | 1.00 | 6 |
| Pre uCB Cell | SQN | -0.08 (-0.11, -0.05) | 0.32 | 2 | -0.09 (-0.12, -0.05) | 0.12 | 1 |
| Post NoCB Unadj | BMIQ | -0.14 (-0.21, -0.07) | 1.00 | 6 | -0.12 (-0.20, -0.04) | 1.00 | 496 |
| Post NoCB Batch | BMIQ | -0.15 (-0.22, -0.09) | 0.65 | 3 | -0.11 (-0.19, -0.03) | 0.68 | 5031 |
| Post NoCB Batch+Cell | BMIQ | -0.16 (-0.22, -0.09) | 0.58 | 1 | -0.15 (-0.22, -0.08) | 0.97 | 18 |
| Post sCB Unadj | BMIQ | -0.15 (-0.21, -0.09) | 0.06 | 2 | -0.11 (-0.18, -0.04) | 0.32 | 3829 |
| Post sCB Cell | BMIQ | -0.16 (-0.22, -0.10) | 0.02 | 1 | -0.14 (-0.20, -0.08) | 0.18 | 34 |
| Post uCB Unadj | BMIQ | -0.14 (-0.20, -0.08) | 0.45 | 2 | -0.11 (-0.18, -0.03) | 0.63 | 2757 |
| Post uCB Cell | BMIQ | -0.15 (-0.21, -0.09) | 0.19 | 1 | -0.14 (-0.20, -0.07) | 0.78 | 14 |
| Pre NoCB Unadj | BMIQ | -0.15 (-0.22, -0.07) | 1.00 | 3 | -0.12 (-0.20, -0.04) | 1.00 | 470 |
| Pre NoCB Batch | BMIQ | -0.16 (-0.23, -0.09) | 0.66 | 3 | -0.11 (-0.19, -0.03) | 0.66 | 4354 |
| Pre NoCB Batch+Cell | BMIQ | -0.17 (-0.24, -0.10) | 0.53 | 1 | -0.15 (-0.22, -0.08) | 0.99 | 14 |
| Pre sCB Unadj | BMIQ | -0.16 (-0.22, -0.10) | 0.05 | 2 | -0.11 (-0.18, -0.04) | 0.31 | 3572 |
| **Pre sCB Cell** | **BMIQ** | **-0.17 (-0.23, -0.11)** | **0.02** | 1 | -0.14 (-0.20, -0.08) | 0.17 | 38 |
| Pre uCB Unadj | BMIQ | -0.15 (-0.21, -0.09) | 0.44 | 1 | -0.11 (-0.18, -0.03) | 0.61 | 2655 |

| | | | | | | | |
|---|---|---|---|---|---|---|---|
| Pre uCB Cell | BMIQ | -0.16 (-0.22, -0.10) | 0.15 | 1 | -0.14 (-0.20, -0.07) | 0.70 | 13 |
| Post NoCB Unadj | SWAN | -0.10 (-0.15, -0.05) | 1.00 | 5 | -0.06 (-0.12, -0.01) | 1.00 | 3127 |
| Post NoCB Batch | SWAN | -0.11 (-0.15, -0.06) | 0.83 | 4 | -0.05 (-0.11, 0.00) | 0.83 | 24600 |
| Post NoCB Batch+Cell | SWAN | -0.11 (-0.16, -0.06) | 1.00 | 1 | -0.08 (-0.12, -0.04) | 1.00 | 12 |
| Post sCB Unadj | SWAN | -0.10 (-0.15, -0.06) | 0.13 | 5 | -0.05 (-0.10, -0.01) | 0.58 | 19768 |
| Post sCB Cell | SWAN | -0.11 (-0.15, -0.07) | 0.10 | 1 | -0.08 (-0.11, -0.04) | 0.53 | 59 |
| Post uCB Unadj | SWAN | -0.10 (-0.14, -0.05) | 0.79 | 5 | -0.05 (-0.10, -0.01) | 0.79 | 13319 |
| Post uCB Cell | SWAN | -0.10 (-0.15, -0.06) | 0.67 | 1 | -0.08 (-0.11, -0.04) | 1.00 | 22 |
| Pre NoCB Unadj | SWAN | -0.10 (-0.15, -0.05) | 1.00 | 5 | -0.06 (-0.12, -0.01) | 1.00 | 3472 |
| Pre NoCB Batch | SWAN | -0.11 (-0.15, -0.06) | 0.83 | 4 | -0.05 (-0.11, 0.00) | 0.83 | 24600 |
| Pre NoCB Batch+Cell | SWAN | -0.11 (-0.15, -0.06) | 1.00 | 1 | -0.08 (-0.12, -0.04) | 1.00 | 12 |
| Pre sCB Unadj | SWAN | -0.10 (-0.14, -0.06) | 0.12 | 5 | -0.05 (-0.10, -0.01) | 0.57 | 21423 |
| Pre sCB Cell | SWAN | -0.11 (-0.15, -0.07) | 0.09 | 1 | -0.08 (-0.11, -0.04) | 0.51 | 65 |
| Pre uCB Unadj | SWAN | -0.09 (-0.13, -0.05) | 0.75 | 5 | -0.05 (-0.10, -0.00) | 0.79 | 14563 |
| Pre uCB Cell | SWAN | -0.10 (-0.14, -0.06) | 0.65 | 1 | -0.08 (-0.11, -0.04) | 1.00 | 24 |

### Table S4: BMI in Standard Care

As there were a large number of significant probes associated with this effect, the 3 probes with the smallest p values (<0.001) were reported here.

| mod | norm | cg06103657 (chr12 PKP2) | | | cg08280341 (chr2 unnamed) | | | cg24630195 (chr5 IRX2) | | |
|---|---|---|---|---|---|---|---|---|---|---|
| | | logFC (95% CI)) | adj.P.Val. | rank. | logFC (95% CI)) | adj.P.Val. | rank. | logFC (95% CI)) | adj.P.Val. | rank. |
| Post NoCB Unadj | SQN | -0.02 (-0.04, -0.00) | 1.00 | 6979 | -0.02 (-0.05, 0.01) | 1.00 | 98728 | 0.02 (-0.01, 0.05) | 1.00 | 112240 |
| Post NoCB Batch | SQN | -0.02 (-0.04, -0.00) | 0.98 | 7310 | -0.01 (-0.04, 0.02) | 0.99 | 289710 | 0.01 (-0.02, 0.05) | 0.98 | 166740 |
| Post NoCB Batch+Cell | SQN | -0.02 (-0.03, -0.00) | 0.78 | 11521 | -0.00 (-0.03, 0.03) | 0.99 | 383606 | 0.01 (-0.02, 0.04) | 0.97 | 271007 |
| Post sCB Unadj | SQN | -0.01 (-0.02, 0.01) | 0.87 | 137943 | -0.01 (-0.03, 0.02) | 0.94 | 258940 | 0.02 (-0.01, 0.04) | 0.84 | 108606 |
| Post sCB Cell | SQN | -0.01 (-0.02, 0.01) | 0.86 | 144436 | -0.00 (-0.03, 0.02) | 0.97 | 320201 | 0.01 (-0.01, 0.04) | 0.89 | 186715 |
| Post uCB Unadj | SQN | -0.01 (-0.02, 0.01) | 1.00 | 171985 | -0.01 (-0.03, 0.02) | 1.00 | 250938 | 0.01 (-0.01, 0.04) | 1.00 | 111003 |
| Post uCB Cell | SQN | -0.00 (-0.02, 0.01) | 1.00 | 182873 | -0.00 (-0.03, 0.02) | 1.00 | 320911 | 0.01 (-0.02, 0.03) | 1.00 | 203580 |
| Pre NoCB Unadj | SQN | -0.02 (-0.03, -0.00) | 1.00 | 14167 | -0.02 (-0.05, 0.01) | 1.00 | 87489 | 0.02 (-0.01, 0.05) | 1.00 | 107615 |
| Pre NoCB Batch | SQN | -0.02 (-0.03, -0.00) | 0.98 | 13475 | -0.01 (-0.04, 0.02) | 0.99 | 264770 | 0.01 (-0.02, 0.05) | 0.98 | 162374 |
| Pre NoCB Batch+Cell | SQN | -0.02 (-0.03, -0.00) | 0.80 | 20236 | -0.00 (-0.03, 0.03) | 0.99 | 355713 | 0.01 (-0.02, 0.04) | 0.97 | 266375 |
| Pre sCB Unadj | SQN | -0.01 (-0.02, 0.01) | 0.88 | 153268 | -0.01 (-0.03, 0.02) | 0.94 | 250386 | 0.02 (-0.01, 0.04) | 0.84 | 105081 |
| Pre sCB Cell | SQN | -0.01 (-0.02, 0.01) | 0.87 | 158565 | -0.01 (-0.03, 0.02) | 0.96 | 309195 | 0.01 (-0.01, 0.04) | 0.89 | 181387 |
| Pre uCB Unadj | SQN | -0.01 (-0.02, 0.01) | 1.00 | 187272 | -0.01 (-0.03, 0.02) | 1.00 | 245327 | 0.02 (-0.01, 0.04) | 1.00 | 107591 |

| | | | | | | | | | | |
|---|---|---|---|---|---|---|---|---|---|---|
| Pre uCB Cell | SQN | -0.00 (-0.02, 0.01) | 1.00 | 195809 | -0.00 (-0.03, 0.02) | 1.00 | 312387 | 0.01 (-0.02, 0.03) | 1.00 | 197979 |
| Post NoCB Unadj | **BMIQ** | **-0.61 (-0.81, -0.40)** | **0.00** | **2** | **-0.61 (-0.81, -0.41)** | **0.00** | **1** | **-0.56 (-0.75, -0.37)** | **0.00** | **3** |
| Post NoCB Batch | **BMIQ** | **-0.68 (-0.89, -0.46)** | **0.00** | **1** | **-0.65 (-0.86, -0.44)** | **0.00** | **3** | **-0.62 (-0.82, -0.42)** | **0.00** | **2** |
| Post NoCB Batch+Cell | **BMIQ** | **-0.67 (-0.89, -0.45)** | **0.00** | **1** | **-0.63 (-0.85, -0.42)** | **0.00** | **3** | **-0.62 (-0.82, -0.42)** | **0.00** | **2** |
| Post sCB Unadj | BMIQ | 0.07 (0.00, 0.14) | 0.45 | 46741 | 0.07 (-0.00, 0.14) | 0.48 | 55933 | 0.07 (0.00, 0.14) | 0.44 | 43865 |
| Post sCB Cell | BMIQ | 0.08 (0.02, 0.14) | 0.35 | 17361 | 0.08 (0.01, 0.15) | 0.39 | 25048 | 0.07 (-0.00, 0.13) | 0.49 | 50121 |
| Post uCB Unadj | BMIQ | -0.11 (-0.17, -0.05) | 0.74 | 86 | -0.09 (-0.16, -0.03) | 0.75 | 1558 | -0.09 (-0.15, -0.03) | 0.75 | 2157 |
| Post uCB Cell | BMIQ | -0.10 (-0.15, -0.05) | 0.70 | 86 | -0.08 (-0.14, -0.02) | 0.76 | 4050 | -0.09 (-0.15, -0.03) | 0.75 | 1247 |
| Pre NoCB Unadj | **BMIQ** | **-0.66 (-0.88, -0.44)** | **0.00** | **2** | **-0.65 (-0.87, -0.44)** | **0.00** | **1** | **-0.61 (-0.81, -0.40)** | **0.00** | 3 |
| Pre NoCB Batch | **BMIQ** | **-0.73 (-0.97, -0.50)** | **0.00** | **1** | **-0.70 (-0.93, -0.47)** | **0.00** | **3** | **-0.67 (-0.89, -0.45)** | **0.00** | 2 |
| Pre NoCB Batch+Cell | **BMIQ** | **-0.72 (-0.96, -0.49)** | **0.00** | **1** | **-0.69 (-0.92, -0.46)** | **0.00** | **3** | **-0.67 (-0.89, -0.45)** | **0.00** | 2 |
| Pre sCB Unadj | BMIQ | 0.07 (0.00, 0.15) | 0.39 | 46303 | 0.08 (0.00, 0.15) | 0.40 | 50061 | 0.08 (0.01, 0.15) | 0.35 | 34922 |
| Pre sCB Cell | BMIQ | 0.09 (0.02, 0.15) | 0.27 | 19486 | 0.09 (0.02, 0.16) | 0.30 | 24249 | 0.08 (0.01, 0.15) | 0.37 | 40413 |
| Pre uCB Unadj | BMIQ | -0.12 (-0.18, -0.06) | 0.49 | 75 | -0.10 (-0.17, -0.04) | 0.55 | 1360 | -0.10 (-0.15, -0.04) | 0.55 | 1293 |
| Pre uCB Cell | BMIQ | -0.11 (-0.16, -0.05) | 0.41 | 89 | -0.09 (-0.15, -0.03) | 0.53 | 3622 | -0.10 (-0.16, -0.04) | 0.48 | 859 |
| Post NoCB Unadj | SWAN | -0.03 (-0.06, -0.00) | 0.82 | 14313 | -0.03 (-0.06, 0.01) | 0.95 | 74767 | 0.01 (-0.02, 0.05) | 0.99 | 187411 |
| Post NoCB Batch | SWAN | -0.04 (-0.07, -0.02) | 0.63 | 1985 | -0.02 (-0.06, 0.02) | 0.77 | 149439 | 0.00 (-0.03, 0.03) | 0.98 | 398746 |
| Post NoCB Batch+Cell | SWAN | -0.03 (-0.06, -0.01) | 0.53 | 1543 | -0.01 (-0.04, 0.02) | 0.93 | 277326 | 0.00 (-0.03, 0.03) | 0.99 | 406007 |
| Post sCB Unadj | SWAN | -0.02 (-0.05, -0.00) | 0.50 | 40012 | -0.02 (-0.05, 0.01) | 0.69 | 151497 | 0.01 (-0.02, 0.03) | 0.91 | 324957 |
| Post sCB Cell | SWAN | -0.02 (-0.04, 0.00) | 0.54 | 44100 | -0.01 (-0.04, 0.02) | 0.85 | 229123 | 0.00 (-0.02, 0.03) | 0.96 | 352754 |
| Post uCB Unadj | SWAN | -0.02 (-0.04, 0.01) | 0.88 | 58712 | -0.02 (-0.05, 0.01) | 0.89 | 149240 | 0.01 (-0.02, 0.03) | 0.96 | 313817 |
| Post uCB Cell | SWAN | -0.01 (-0.03, 0.01) | 0.93 | 81657 | -0.01 (-0.04, 0.02) | 0.98 | 244896 | 0.00 (-0.02, 0.03) | 0.99 | 346014 |
| Pre NoCB Unadj | SWAN | -0.03 (-0.06, -0.00) | 0.81 | 14996 | -0.03 (-0.06, 0.01) | 0.94 | 79456 | 0.01 (-0.02, 0.05) | 0.98 | 191950 |
| Pre NoCB Batch | SWAN | -0.04 (-0.07, -0.02) | 0.63 | 1985 | -0.02 (-0.06, 0.02) | 0.77 | 149439 | 0.00 (-0.03, 0.03) | 0.98 | 398746 |
| Pre NoCB Batch+Cell | SWAN | -0.03 (-0.06, -0.01) | 0.53 | 2147 | -0.01 (-0.04, 0.02) | 0.94 | 283064 | 0.00 (-0.03, 0.03) | 1.00 | 412966 |
| Pre sCB Unadj | SWAN | -0.02 (-0.05, -0.00) | 0.50 | 41187 | -0.02 (-0.05, 0.01) | 0.69 | 149901 | 0.01 (-0.02, 0.03) | 0.92 | 333111 |
| Pre sCB Cell | SWAN | -0.02 (-0.04, 0.00) | 0.54 | 46002 | -0.01 (-0.04, 0.02) | 0.85 | 224799 | 0.00 (-0.02, 0.03) | 0.96 | 360925 |

| | | | | | | | | | | |
|---|---|---|---|---|---|---|---|---|---|---|
| Pre uCB Unadj | SWAN | -0.02 (-0.04, 0.01) | 0.88 | 59824 | -0.02 (-0.05, 0.02) | 0.89 | 148048 | 0.01 (-0.02, 0.03) | 0.96 | 321848 |
| Pre uCB Cell | SWAN | -0.01 (-0.03, 0.01) | 0.93 | 83966 | -0.01 (-0.04, 0.02) | 0.97 | 240839 | 0.00 (-0.03, 0.03) | 0.99 | 354388 |

### Table S5: BMI in Lifestyle Advice

| mod | | cg07823293 (chr11 TBRG1) | | | cg02727674 (chr12 FGD6; VEZT) | | | cg27347003 (chr16 ATP2A1) | | |
|---|---|---|---|---|---|---|---|---|---|---|
| | norm | logFC (95% CI) | adj.P.Val. | rank. | logFC (95% CI) | adj.P.Val. | rank. | logFC (95% CI) | adj.P.Val. | rank. |
| Post NoCB Unadj | SQN | 0.08 (0.05, 0.12) | 0.87 | 3 | 0.04 (0.02, 0.05) | 1.00 | 16 | -0.02 (-0.05, 0.01) | 1.00 | 112808 |
| Post NoCB Batch | SQN | 0.09 (0.05, 0.12) | 0.23 | 2 | 0.04 (0.02, 0.05) | 0.95 | 39 | -0.01 (-0.04, 0.02) | 0.99 | 187420 |
| Post NoCB Batch+Cell | SQN | 0.09 (0.05, 0.12) | 0.40 | 1 | 0.04 (0.02, 0.05) | 0.85 | 42 | -0.01 (-0.04, 0.02) | 1.00 | 154477 |
| Post sCB Unadj | SQN | 0.08 (0.05, 0.11) | 0.05 | 1 | 0.01 (-0.00, 0.03) | 0.76 | 53236 | -0.01 (-0.04, 0.01) | 0.87 | 134114 |
| Post sCB Cell | **SQN** | **0.08 (0.05, 0.12)** | **0.03** | **1** | 0.01 (-0.00, 0.03) | 0.66 | 45473 | -0.02 (-0.04, 0.01) | 0.79 | 111149 |
| Post uCB Unadj | SQN | 0.08 (0.05, 0.11) | 0.43 | 1 | 0.01 (-0.00, 0.03) | 1.00 | 46494 | -0.01 (-0.04, 0.01) | 1.00 | 130173 |
| Post uCB Cell | SQN | 0.08 (0.05, 0.11) | 0.29 | 1 | 0.01 (-0.00, 0.03) | 0.91 | 38174 | -0.02 (-0.04, 0.01) | 0.94 | 104444 |
| Pre NoCB Unadj | SQN | 0.08 (0.05, 0.12) | 0.83 | 3 | 0.03 (0.02, 0.05) | 1.00 | 38 | -0.02 (-0.05, 0.01) | 1.00 | 109076 |
| Pre NoCB Batch | SQN | 0.09 (0.05, 0.12) | 0.23 | 2 | 0.03 (0.02, 0.05) | 1.00 | 84 | -0.01 (-0.04, 0.02) | 1.00 | 184438 |
| Pre NoCB Batch+Cell | SQN | 0.09 (0.05, 0.13) | 0.38 | 1 | 0.03 (0.02, 0.05) | 0.86 | 86 | -0.01 (-0.04, 0.02) | 1.00 | 155581 |
| Pre sCB Unadj | SQN | 0.09 (0.05, 0.12) | 0.05 | 1 | 0.01 (-0.00, 0.03) | 0.75 | 48166 | -0.01 (-0.04, 0.01) | 0.87 | 133875 |
| Pre sCB Cell | **SQN** | **0.09 (0.06, 0.12)** | **0.03** | 1 | 0.01 (-0.00, 0.03) | 0.65 | 40295 | -0.02 (-0.04, 0.01) | 0.79 | 112662 |
| Pre uCB Unadj | SQN | 0.08 (0.05, 0.11) | 0.43 | 1 | 0.01 (-0.00, 0.02) | 1.00 | 49799 | -0.01 (-0.04, 0.01) | 1.00 | 130388 |
| Pre uCB Cell | SQN | 0.08 (0.05, 0.11) | 0.29 | 1 | 0.01 (-0.00, 0.02) | 0.91 | 40186 | -0.02 (-0.04, 0.01) | 0.94 | 106674 |
| Post NoCB Unadj | BMIQ | 0.07 (0.03, 0.10) | 1.00 | 27 | **-0.48 (-0.66, -0.30)** | **0.04** | **2** | **-0.55 (-0.76, -0.35)** | **0.04** | **1** |
| Post NoCB Batch | BMIQ | 0.07 (0.04, 0.10) | 0.97 | 9 | -0.50 (-0.69, -0.31) | 0.05 | 2 | -0.56 (-0.77, -0.35) | 0.05 | 1 |
| Post NoCB Batch+Cell | BMIQ | 0.07 (0.04, 0.10) | 0.81 | 8 | -0.46 (-0.65, -0.28) | 0.13 | 4 | -0.53 (-0.74, -0.32) | 0.11 | 1 |
| Post sCB Unadj | BMIQ | 0.06 (0.04, 0.09) | 0.27 | 7 | 0.10 (0.04, 0.15) | 0.45 | 346 | 0.09 (0.02, 0.16) | 0.66 | 5952 |
| Post sCB Cell | BMIQ | 0.06 (0.04, 0.09) | 0.16 | 13 | 0.10 (0.05, 0.15) | 0.27 | 215 | 0.09 (0.02, 0.16) | 0.52 | 8199 |
| Post uCB Unadj | BMIQ | 0.06 (0.03, 0.09) | 1.00 | 5 | -0.07 (-0.11, -0.02) | 1.00 | 723 | -0.09 (-0.15, -0.03) | 1.00 | 1114 |
| Post uCB Cell | BMIQ | 0.06 (0.03, 0.09) | 1.00 | 8 | -0.06 (-0.11, -0.02) | 1.00 | 2242 | -0.09 (-0.15, -0.03) | 1.00 | 1297 |
| Pre NoCB Unadj | BMIQ | 0.07 (0.03, 0.10) | 1.00 | 24 | -0.50 (-0.69, -0.31) | 0.08 | 2 | -0.58 (-0.79, -0.37) | 0.05 | 1 |

| | | | | | | | | | | |
|---|---|---|---|---|---|---|---|---|---|---|
| Pre NoCB Batch | BMIQ | 0.07 (0.04, 0.10) | 1.00 | 10 | -0.52 (-0.71, -0.32) | 0.09 | 2 | -0.59 (-0.80, -0.37) | 0.08 | 1 |
| Pre NoCB Batch+Cell | BMIQ | 0.07 (0.04, 0.10) | 0.90 | 8 | -0.48 (-0.67, -0.28) | 0.22 | 4 | -0.55 (-0.77, -0.34) | 0.11 | 2 |
| Pre sCB Unadj | BMIQ | 0.06 (0.04, 0.09) | 0.34 | 5 | 0.12 (0.06, 0.17) | 0.45 | 30 | 0.10 (0.03, 0.17) | 0.73 | 3459 |
| Pre sCB Cell | BMIQ | 0.06 (0.04, 0.09) | 0.21 | 9 | 0.12 (0.07, 0.18) | 0.24 | 19 | 0.10 (0.03, 0.17) | 0.57 | 4795 |
| Pre uCB Unadj | BMIQ | 0.06 (0.03, 0.09) | 1.00 | 6 | -0.05 (-0.10, -0.01) | 1.00 | 6529 | -0.09 (-0.15, -0.03) | 1.00 | 982 |
| Pre uCB Cell | BMIQ | 0.06 (0.03, 0.09) | 1.00 | 7 | -0.05 (-0.09, -0.00) | 1.00 | 13512 | -0.09 (-0.15, -0.03) | 1.00 | 1082 |
| Post NoCB Unadj | SWAN | 0.06 (0.03, 0.09) | 1.00 | 21 | 0.04 (0.01, 0.06) | 1.00 | 294 | -0.03 (-0.07, 0.02) | 1.00 | 51398 |
| Post NoCB Batch | SWAN | 0.06 (0.03, 0.09) | 1.00 | 18 | 0.04 (0.01, 0.06) | 1.00 | 301 | -0.02 (-0.06, 0.01) | 1.00 | 79783 |
| Post NoCB Batch+Cell | SWAN | 0.06 (0.03, 0.09) | 0.97 | 39 | 0.03 (0.01, 0.06) | 0.97 | 1080 | -0.03 (-0.06, 0.01) | 0.97 | 47271 |
| Post sCB Unadj | SWAN | 0.05 (0.03, 0.08) | 0.53 | 36 | 0.01 (-0.01, 0.03) | 0.96 | 73778 | -0.02 (-0.05, 0.01) | 0.96 | 75198 |
| Post sCB Cell | SWAN | 0.05 (0.03, 0.08) | 0.29 | 67 | 0.01 (-0.01, 0.03) | 0.83 | 90135 | -0.02 (-0.05, 0.01) | 0.78 | 58949 |
| Post uCB Unadj | SWAN | 0.05 (0.02, 0.08) | 1.00 | 29 | 0.01 (-0.01, 0.03) | 1.00 | 71666 | -0.02 (-0.05, 0.01) | 1.00 | 79453 |
| Post uCB Cell | SWAN | 0.05 (0.02, 0.08) | 1.00 | 53 | 0.01 (-0.01, 0.03) | 1.00 | 86737 | -0.02 (-0.05, 0.01) | 1.00 | 59519 |
| Pre NoCB Unadj | SWAN | 0.06 (0.03, 0.09) | 1.00 | 19 | 0.04 (0.01, 0.06) | 1.00 | 333 | -0.03 (-0.07, 0.02) | 1.00 | 53126 |
| Pre NoCB Batch | SWAN | 0.06 (0.03, 0.09) | 1.00 | 18 | 0.04 (0.01, 0.06) | 1.00 | 301 | -0.02 (-0.06, 0.01) | 1.00 | 79783 |
| Pre NoCB Batch+Cell | SWAN | 0.06 (0.03, 0.09) | 0.97 | 42 | 0.04 (0.01, 0.06) | 0.98 | 1288 | -0.03 (-0.06, 0.01) | 0.98 | 49671 |
| Pre sCB Unadj | SWAN | 0.05 (0.03, 0.08) | 0.51 | 36 | 0.01 (-0.01, 0.03) | 0.96 | 76313 | -0.02 (-0.05, 0.01) | 0.96 | 71517 |
| Pre sCB Cell | SWAN | 0.05 (0.03, 0.08) | 0.28 | 72 | 0.01 (-0.01, 0.03) | 0.83 | 92257 | -0.02 (-0.05, 0.00) | 0.77 | 55607 |
| Pre uCB Unadj | SWAN | 0.05 (0.02, 0.08) | 1.00 | 30 | 0.01 (-0.01, 0.03) | 1.00 | 76464 | -0.02 (-0.05, 0.01) | 1.00 | 74464 |
| Pre uCB Cell | SWAN | 0.05 (0.02, 0.08) | 1.00 | 56 | 0.01 (-0.01, 0.03) | 1.00 | 91323 | -0.02 (-0.05, 0.01) | 1.00 | 55442 |

### Tables: S6-S8 Fake Group Effects

Note that there is no table for Tortoiseshell in Short-Haired as there were no DMPs for this effect in any of the models.

#### Table S6: Tortoiseshell in Long-Haired

| | | cg05704183 (chr2 unnamed) | | | cg19644590 (chr19 unnamed) | | | cg09783457 (chr4 unnamed) | | |
|---|---|---|---|---|---|---|---|---|---|---|
| mod | norm | logFC (95% CI) | adj.P.Val. | rank | logFC (95% CI) | adj.P.Val. | rank | logFC (95% CI) | adj.P.Val. | rank |
| Post NoCB Unadj | SQN | 0.12 (0.07, 0.17) | 0.66 | 3 | -0.10 (-0.14, -0.05) | 0.66 | 9 | 0.09 (0.04, 0.14) | 0.66 | 59 |
| Post NoCB Batch | SQN | 0.13 (0.08, 0.18) | 0.27 | 2 | -0.10 (-0.14, -0.05) | 0.27 | 10 | 0.11 (0.06, 0.16) | 0.27 | 4 |

| | | | | | | | | | | |
|---|---|---|---|---|---|---|---|---|---|---|
| Post NoCB Batch+Cell | SQN | 0.11 (0.06, 0.16) | 0.70 | 4 | -0.08 (-0.12, -0.04) | 0.70 | 10 | 0.10 (0.05, 0.14) | 0.70 | 20 |
| Post sCB Unadj | SQN | **0.13 (0.08, 0.17)** | **0.03** | **2** | **-0.10 (-0.14, -0.06)** | **0.03** | **1** | **0.11 (0.07, 0.15)** | **0.03** | **3** |
| Post sCB Cell | SQN | 0.11 (0.07, 0.16) | 0.05 | 3 | -0.09 (-0.12, -0.06) | 0.05 | 1 | 0.10 (0.06, 0.14) | 0.05 | 7 |
| Post uCB Unadj | SQN | 0.11 (0.06, 0.16) | 0.34 | 3 | -0.09 (-0.12, -0.05) | 0.34 | 1 | 0.10 (0.05, 0.14) | 0.34 | 7 |
| Post uCB Cell | SQN | 0.10 (0.05, 0.14) | 0.69 | 8 | -0.08 (-0.11, -0.04) | 0.69 | 2 | 0.09 (0.05, 0.13) | 0.69 | 14 |
| Pre NoCB Unadj | SQN | 0.12 (0.07, 0.18) | 0.66 | 3 | -0.09 (-0.14, -0.05) | 0.66 | 8 | 0.10 (0.05, 0.14) | 0.66 | 54 |
| Pre NoCB Batch | SQN | 0.13 (0.08, 0.19) | 0.27 | 2 | -0.09 (-0.13, -0.05) | 0.27 | 11 | 0.11 (0.07, 0.16) | 0.27 | 4 |
| Pre NoCB Batch+Cell | SQN | 0.11 (0.06, 0.17) | 0.68 | 4 | -0.08 (-0.12, -0.04) | 0.68 | 11 | 0.10 (0.05, 0.15) | 0.68 | 20 |
| Pre sCB Unadj | SQN | **0.13 (0.08, 0.17)** | **0.03** | **3** | **-0.10 (-0.13, -0.06)** | **0.03** | **1** | **0.11 (0.07, 0.16)** | **0.03** | **2** |
| Pre sCB Cell | SQN | 0.11 (0.07, 0.16) | 0.05 | 3 | -0.09 (-0.12, -0.06) | 0.05 | 1 | 0.10 (0.06, 0.14) | 0.05 | 7 |
| Pre uCB Unadj | SQN | 0.11 (0.06, 0.16) | 0.35 | 3 | -0.08 (-0.12, -0.05) | 0.35 | 1 | 0.10 (0.06, 0.14) | 0.35 | 6 |
| Pre uCB Cell | SQN | 0.10 (0.05, 0.14) | 0.69 | 8 | -0.08 (-0.11, -0.04) | 0.69 | 2 | 0.09 (0.05, 0.13) | 0.69 | 13 |
| Post NoCB Unadj | BMIQ | 0.22 (0.12, 0.32) | 1.00 | 1 | -0.12 (-0.20, -0.04) | 1.00 | 237 | 0.14 (0.06, 0.22) | 1.00 | 62 |
| Post NoCB Batch | BMIQ | 0.21 (0.11, 0.31) | 0.60 | 11 | -0.13 (-0.20, -0.05) | 0.60 | 530 | 0.15 (0.07, 0.23) | 0.60 | 306 |
| Post NoCB Batch+Cell | BMIQ | 0.21 (0.11, 0.30) | 0.93 | 6 | -0.10 (-0.17, -0.03) | 0.93 | 1479 | 0.16 (0.08, 0.24) | 0.93 | 27 |
| Post sCB Unadj | BMIQ | 0.21 (0.12, 0.30) | 0.15 | 7 | -0.13 (-0.20, -0.07) | 0.16 | 166 | 0.15 (0.08, 0.22) | 0.16 | 115 |
| Post sCB Cell | BMIQ | 0.21 (0.12, 0.29) | 0.22 | 2 | -0.11 (-0.17, -0.05) | 0.35 | 361 | 0.16 (0.09, 0.23) | 0.23 | 14 |
| Post uCB Unadj | BMIQ | 0.18 (0.10, 0.27) | 0.81 | 9 | -0.11 (-0.18, -0.05) | 0.81 | 195 | 0.14 (0.06, 0.21) | 0.81 | 104 |
| Post uCB Cell | BMIQ | 0.18 (0.10, 0.27) | 1.00 | 2 | -0.10 (-0.16, -0.03) | 1.00 | 510 | 0.14 (0.07, 0.21) | 1.00 | 10 |
| Pre NoCB Unadj | BMIQ | 0.23 (0.13, 0.33) | 1.00 | 1 | -0.13 (-0.21, -0.05) | 1.00 | 263 | 0.15 (0.06, 0.23) | 1.00 | 75 |
| Pre NoCB Batch | BMIQ | 0.22 (0.12, 0.32) | 0.63 | 12 | -0.13 (-0.21, -0.06) | 0.63 | 382 | 0.15 (0.07, 0.24) | 0.63 | 278 |
| Pre NoCB Batch+Cell | BMIQ | 0.21 (0.12, 0.31) | 1.00 | 5 | -0.11 (-0.18, -0.04) | 1.00 | 905 | 0.16 (0.08, 0.24) | 1.00 | 33 |
| Pre sCB Unadj | BMIQ | 0.22 (0.13, 0.31) | 0.12 | 8 | -0.14 (-0.20, -0.07) | 0.16 | 117 | 0.16 (0.08, 0.23) | 0.16 | 125 |
| Pre sCB Cell | BMIQ | 0.21 (0.13, 0.30) | 0.20 | 3 | -0.12 (-0.18, -0.06) | 0.35 | 248 | 0.16 (0.09, 0.23) | 0.26 | 16 |
| Pre uCB Unadj | BMIQ | 0.19 (0.10, 0.28) | 0.81 | 10 | -0.12 (-0.18, -0.05) | 0.81 | 151 | 0.14 (0.06, 0.21) | 0.81 | 101 |
| Pre uCB Cell | BMIQ | 0.19 (0.10, 0.27) | 1.00 | 2 | -0.10 (-0.16, -0.04) | 1.00 | 379 | 0.14 (0.07, 0.22) | 1.00 | 13 |
| Post NoCB Unadj | SWAN | 0.14 (0.07, 0.21) | 1.00 | 2 | -0.09 (-0.15, -0.02) | 1.00 | 2122 | 0.10 (0.03, 0.17) | 1.00 | 591 |

| | | | | | | | | | | |
|---|---|---|---|---|---|---|---|---|---|---|
| Post NoCB Batch | SWAN | 0.12 (0.05, 0.19) | 0.36 | 967 | -0.10 (-0.17, -0.04) | 0.36 | 1640 | 0.09 (0.02, 0.16) | 0.40 | 11263 |
| Post NoCB Batch+Cell | SWAN | 0.13 (0.07, 0.20) | 0.66 | 13 | -0.07 (-0.12, -0.02) | 0.66 | 4587 | 0.11 (0.05, 0.17) | 0.66 | 87 |
| Post sCB Unadj | SWAN | 0.12 (0.06, 0.18) | 0.12 | 587 | -0.11 (-0.16, -0.05) | 0.13 | 734 | 0.09 (0.03, 0.15) | 0.19 | 7453 |
| Post sCB Cell | SWAN | 0.13 (0.07, 0.19) | 0.23 | 11 | -0.08 (-0.12, -0.03) | 0.28 | 1204 | 0.10 (0.05, 0.15) | 0.24 | 157 |
| Post uCB Unadj | SWAN | 0.10 (0.04, 0.17) | 0.53 | 621 | -0.09 (-0.15, -0.04) | 0.53 | 893 | 0.08 (0.02, 0.14) | 0.54 | 5586 |
| Post uCB Cell | SWAN | 0.12 (0.06, 0.17) | 0.93 | 8 | -0.06 (-0.11, -0.02) | 0.93 | 1999 | 0.10 (0.04, 0.15) | 0.93 | 57 |
| Pre NoCB Unadj | SWAN | 0.15 (0.08, 0.22) | 1.00 | 2 | -0.09 (-0.16, -0.02) | 1.00 | 1607 | 0.10 (0.03, 0.18) | 1.00 | 541 |
| Pre NoCB Batch | SWAN | 0.13 (0.06, 0.20) | 0.34 | 647 | -0.11 (-0.18, -0.05) | 0.34 | 1150 | 0.10 (0.03, 0.17) | 0.37 | 9027 |
| Pre NoCB Batch+Cell | SWAN | 0.14 (0.08, 0.21) | 0.64 | 10 | -0.07 (-0.12, -0.02) | 0.64 | 3157 | 0.11 (0.06, 0.17) | 0.64 | 71 |
| Pre sCB Unadj | SWAN | 0.13 (0.06, 0.19) | 0.11 | 365 | -0.11 (-0.17, -0.05) | 0.11 | 520 | 0.10 (0.03, 0.16) | 0.17 | 5889 |
| Pre sCB Cell | SWAN | 0.14 (0.08, 0.19) | 0.17 | 7 | -0.08 (-0.13, -0.04) | 0.26 | 795 | 0.11 (0.06, 0.16) | 0.23 | 116 |
| Pre uCB Unadj | SWAN | 0.11 (0.05, 0.17) | 0.51 | 400 | -0.10 (-0.15, -0.04) | 0.51 | 634 | 0.09 (0.03, 0.15) | 0.51 | 4461 |
| Pre uCB Cell | SWAN | 0.12 (0.06, 0.18) | 0.90 | 5 | -0.07 (-0.11, -0.02) | 0.90 | 1470 | 0.10 (0.05, 0.15) | 0.90 | 48 |

### Table S7: Short-Haired in Tabby

As there were a large number of significant probes, the 2 probes with the smallest p value (<0.005 in at least one model) are reported below.

| | | cg13560436 (chrr5 MIER3) | | | cg07589064 (chr11 unnamed) | | |
|---|---|---|---|---|---|---|---|
| mod | norm | logFC (95% CI) | adj.P.Val. | rank | logFC (95% CI) | adj.P.Val. | rank |
| Post NoCB Unadj | SQN | 0.10 (0.06, 0.15) | 0.65 | 1 | 0.11 (0.06, 0.16) | 0.65 | 6 |
| Post NoCB Batch | SQN | **0.12 (0.08, 0.17)** | **0.03** | **1** | **0.13 (0.08, 0.18)** | **0.03** | **2** |
| Post NoCB Batch+Cell | SQN | 0.10 (0.06, 0.14) | 0.51 | 3 | 0.08 (0.04, 0.12) | 0.81 | 19 |
| Post sCB Unadj | SQN | **0.11 (0.08, 0.15)** | **0.01** | **1** | **0.12 (0.08, 0.16)** | **0.01** | **2** |
| Post sCB Cell | SQN | **0.10 (0.07, 0.14)** | **0.01** | **1** | **0.09 (0.06, 0.13)** | **0.01** | **2** |
| Post uCB Unadj | SQN | 0.10 (0.06, 0.14) | 0.07 | 2 | 0.11 (0.07, 0.16) | 0.07 | 1 |
| Post uCB Cell | SQN | 0.09 (0.06, 0.13) | 0.17 | 1 | 0.08 (0.05, 0.12) | 0.17 | 2 |
| Pre NoCB Unadj | SQN | 0.10 (0.06, 0.15) | 0.64 | 1 | 0.11 (0.06, 0.16) | 0.64 | 6 |
| Pre NoCB Batch | SQN | **0.12 (0.08, 0.17)** | **0.03** | **1** | **0.13 (0.08, 0.18)** | **0.03** | **2** |
| Pre NoCB Batch+Cell | SQN | 0.10 (0.06, 0.14) | 0.51 | 3 | 0.08 (0.04, 0.12) | 0.79 | 19 |
| Pre sCB Unadj | SQN | **0.11 (0.07, 0.15)** | **0.00** | **2** | **0.12 (0.08, 0.16)** | **0.00** | **1** |
| Pre sCB Cell | SQN | **0.10 (0.07, 0.14)** | **0.01** | **1** | **0.09 (0.06, 0.13)** | **0.01** | **2** |
| Pre uCB Unadj | SQN | 0.10 (0.06, 0.14) | 0.07 | 2 | 0.11 (0.07, 0.16) | 0.07 | 1 |
| Pre uCB Cell | SQN | 0.09 (0.06, 0.13) | 0.16 | 2 | 0.08 (0.05, 0.12) | 0.16 | 1 |
| Post NoCB Unadj | BMIQ | 0.10 (0.05, 0.16) | 1.00 | 22 | 0.10 (0.04, 0.16) | 1.00 | 178 |
| Post NoCB Batch | BMIQ | 0.11 (0.06, 0.17) | 0.22 | 110 | 0.10 (0.05, 0.16) | 0.25 | 910 |

| | | | | | | | |
|---|---|---|---|---|---|---|---|
| Post NoCB Batch+Cell | BMIQ | 0.10 (0.05, 0.16) | 1.00 | 34 | 0.10 (0.04, 0.15) | 1.00 | 83 |
| Post sCB Unadj | BMIQ | 0.11 (0.06, 0.16) | 0.07 | 73 | 0.11 (0.06, 0.16) | 0.08 | 150 |
| Post sCB Cell | BMIQ | 0.10 (0.06, 0.15) | 0.20 | 30 | 0.10 (0.05, 0.15) | 0.23 | 87 |
| Post uCB Unadj | BMIQ | 0.10 (0.05, 0.15) | 0.45 | 44 | 0.10 (0.05, 0.15) | 0.45 | 122 |
| Post uCB Cell | BMIQ | 0.10 (0.05, 0.14) | 1.00 | 20 | 0.09 (0.04, 0.14) | 1.00 | 72 |
| Pre NoCB Unadj | BMIQ | 0.11 (0.05, 0.16) | 1.00 | 16 | 0.11 (0.04, 0.17) | 1.00 | 149 |
| Pre NoCB Batch | BMIQ | 0.11 (0.06, 0.17) | 0.22 | 120 | 0.10 (0.04, 0.16) | 0.29 | 2853 |
| Pre NoCB Batch+Cell | BMIQ | 0.10 (0.05, 0.16) | 1.00 | 39 | 0.09 (0.04, 0.15) | 1.00 | 300 |
| Pre sCB Unadj | BMIQ | 0.11 (0.06, 0.16) | 0.07 | 91 | 0.10 (0.05, 0.16) | 0.10 | 619 |
| Pre sCB Cell | BMIQ | 0.10 (0.06, 0.15) | 0.19 | 45 | 0.09 (0.04, 0.14) | 0.30 | 293 |
| Pre uCB Unadj | BMIQ | 0.10 (0.05, 0.15) | 0.42 | 57 | 0.09 (0.04, 0.15) | 0.46 | 420 |
| Pre uCB Cell | BMIQ | 0.09 (0.05, 0.14) | 1.00 | 25 | 0.09 (0.04, 0.14) | 1.00 | 197 |
| Post NoCB Unadj | SWAN | 0.12 (0.06, 0.17) | 1.00 | 1 | 0.07 (0.03, 0.12) | 1.00 | 59 |
| Post NoCB Batch | SWAN | 0.13 (0.08, 0.19) | 0.27 | 1 | 0.08 (0.04, 0.12) | 0.31 | 457 |
| Post NoCB Batch+Cell | SWAN | 0.11 (0.06, 0.16) | 1.00 | 3 | 0.07 (0.03, 0.11) | 1.00 | 108 |
| Post sCB Unadj | SWAN | **0.13 (0.08, 0.17)** | **0.04** | **1** | 0.08 (0.04, 0.11) | 0.10 | 228 |
| Post sCB Cell | SWAN | 0.12 (0.07, 0.16) | 0.09 | 1 | 0.07 (0.03, 0.10) | 0.41 | 160 |
| Post uCB Unadj | SWAN | 0.12 (0.07, 0.16) | 0.46 | 1 | 0.07 (0.03, 0.11) | 0.49 | 215 |
| Post uCB Cell | SWAN | 0.11 (0.06, 0.15) | 1.00 | 1 | 0.06 (0.02, 0.09) | 1.00 | 155 |
| Pre NoCB Unadj | SWAN | 0.12 (0.07, 0.17) | 1.00 | 1 | 0.08 (0.04, 0.12) | 1.00 | 39 |
| Pre NoCB Batch | SWAN | 0.14 (0.08, 0.19) | 0.26 | 1 | 0.09 (0.04, 0.13) | 0.29 | 235 |
| Pre NoCB Batch+Cell | SWAN | 0.12 (0.06, 0.17) | 1.00 | 3 | 0.07 (0.03, 0.11) | 1.00 | 72 |
| Pre sCB Unadj | SWAN | **0.13 (0.08, 0.18)** | **0.04** | **1** | 0.08 (0.04, 0.12) | 0.08 | 114 |
| Pre sCB Cell | SWAN | 0.12 (0.07, 0.16) | 0.07 | 1 | 0.07 (0.04, 0.11) | 0.36 | 80 |
| Pre uCB Unadj | SWAN | 0.12 (0.07, 0.17) | 0.45 | 1 | 0.08 (0.04, 0.11) | 0.45 | 108 |
| Pre uCB Cell | SWAN | 0.11 (0.06, 0.15) | 1.00 | 1 | 0.06 (0.03, 0.10) | 1.00 | 78 |

### Table S8: Short-Haired in Tortoiseshell

| Model | Norm. | cg12432161 (chr3 unnamed) | | |
|---|---|---|---|---|
| | | logFC (95% CI) | adj.P.Val. | rank |
| Post NoCB Unadj | SQN | 0.09 (0.04, 0.13) | 30 | 1.00 |
| Post NoCB Batch | SQN | 0.10 (0.05, 0.14) | 21 | 1.00 |
| Post NoCB Batch+Cell | SQN | 0.09 (0.05, 0.12) | 2 | 0.75 |
| Post sCB Unadj | SQN | 0.10 (0.06, 0.14) | 7 | 0.15 |
| **Post sCB Cell** | **SQN** | **0.09 (0.06, 0.12)** | **1** | **0.03** |
| Post uCB Unadj | SQN | 0.09 (0.05, 0.13) | 10 | 1.00 |
| Post uCB Cell | SQN | 0.08 (0.05, 0.11) | 1 | 1.00 |
| Pre NoCB Unadj | SQN | 0.09 (0.04, 0.13) | 29 | 1.00 |
| Pre NoCB Batch | SQN | 0.09 (0.05, 0.14) | 19 | 1.00 |
| Pre NoCB Batch+Cell | SQN | 0.09 (0.05, 0.12) | 2 | 0.89 |
| Pre sCB Unadj | SQN | 0.10 (0.06, 0.14) | 6 | 0.15 |
| **Pre sCB Cell** | **SQN** | **0.09 (0.06, 0.12)** | **1** | **0.03** |
| Pre uCB Unadj | SQN | 0.09 (0.05, 0.13) | 9 | 1.00 |
| Pre uCB Cell | SQN | 0.08 (0.05, 0.11) | 1 | 1.00 |
| Post NoCB Unadj | BMIQ | 0.16 (0.05, 0.26) | 527 | 1.00 |

| | | | | |
|---|---|---|---|---|
| Post NoCB Batch | BMIQ | 0.20 (0.09, 0.31) | 48 | 1.00 |
| Post NoCB Batch+Cell | BMIQ | 0.16 (0.09, 0.24) | 4 | 1.00 |
| Post sCB Unadj | BMIQ | 0.21 (0.12, 0.31) | 12 | 0.47 |
| Post sCB Cell | BMIQ | 0.19 (0.12, 0.26) | 1 | 0.06 |
| Post uCB Unadj | BMIQ | 0.19 (0.09, 0.28) | 13 | 1.00 |
| Post uCB Cell | BMIQ | 0.16 (0.09, 0.23) | 1 | 1.00 |
| Pre NoCB Unadj | BMIQ | 0.16 (0.05, 0.27) | 361 | 1.00 |
| Pre NoCB Batch | BMIQ | 0.21 (0.10, 0.32) | 37 | 1.00 |
| Pre NoCB Batch+Cell | BMIQ | 0.17 (0.09, 0.24) | 4 | 1.00 |
| Pre sCB Unadj | BMIQ | 0.22 (0.13, 0.32) | 9 | 0.37 |
| **Pre sCB Cell** | **BMIQ** | **0.20 (0.13, 0.27)** | **1** | **0.03** |
| Pre uCB Unadj | BMIQ | 0.20 (0.10, 0.29) | 9 | 1.00 |
| Pre uCB Cell | BMIQ | 0.17 (0.10, 0.24) | 1 | 1.00 |
| Post NoCB Unadj | SWAN | 0.10 (0.01, 0.18) | 2797 | 1.00 |
| Post NoCB Batch | SWAN | 0.13 (0.04, 0.22) | 378 | 1.00 |
| Post NoCB Batch+Cell | SWAN | 0.10 (0.06, 0.15) | 6 | 1.00 |
| Post sCB Unadj | SWAN | 0.13 (0.06, 0.21) | 230 | 0.89 |
| Post sCB Cell | SWAN | 0.11 (0.07, 0.16) | 8 | 0.22 |
| Post uCB Unadj | SWAN | 0.12 (0.04, 0.19) | 212 | 1.00 |
| Post uCB Cell | SWAN | 0.10 (0.05, 0.15) | 9 | 1.00 |
| Pre NoCB Unadj | SWAN | 0.10 (0.02, 0.19) | 2377 | 1.00 |
| Pre NoCB Batch | SWAN | 0.14 (0.05, 0.22) | 318 | 1.00 |
| Pre NoCB Batch+Cell | SWAN | 0.11 (0.06, 0.16) | 6 | 1.00 |
| Pre sCB Unadj | SWAN | 0.14 (0.06, 0.21) | 193 | 0.91 |
| Pre sCB Cell | SWAN | 0.12 (0.07, 0.17) | 6 | 0.23 |
| Pre uCB Unadj | SWAN | 0.12 (0.05, 0.20) | 178 | 1.00 |
| Pre uCB Cell | SWAN | 0.10 (0.05, 0.15) | 7 | 1.00 |
